# Insights into the low-temperature adaptation of an enzyme as studied through ancestral sequence reconstruction

**DOI:** 10.1101/2024.09.05.611558

**Authors:** Shuang Cui, Ryutaro Furukawa, Satoshi Akanuma

**Affiliations:** Faculty of Human Sciences, Waseda University, 2-579-15 Mikajima, Tokorozawa, Saitama 359-1192, Japan

**Keywords:** Activation energy, Ancestral sequence reconstruction, Low-temperature activity, Low-temperature adaptation, Thermostability

## Abstract

For billions of years, enzymes have evolved in response to the changing environments in which their host organisms lived. Various lines of evidence suggest the earliest primitive organisms inhabited high-temperature environments and possessed enzymes adapted to such conditions. Consequently, extant mesophilic and psychrophilic enzymes are believed to have adapted to lower temperatures during the evolutionary process. Herein, we analyzed this low-temperature adaptation using ancestral sequence reconstruction. Previously, we generated the phylogenetic tree of 3-isopropylmalate dehydrogenases (IPMDHs) and reconstructed the sequence of the last bacterial common ancestor. The corresponding ancestral enzyme displayed high thermostability and catalytic activity at elevated temperatures but moderate activity at low temperatures (Furukawa *et al*., *Sci. Rep.* 10, 15493 (2020)). Here, to identify amino acid residues that are responsible for the low-temperature adaptation, we reconstructed and characterized all eleven evolutionary intermediates that sequentially connect the last bacterial common ancestor with extant mesophilic IPMDH from *Escherichia coli*. A remarkable change in catalytic properties, from those suited for high reaction temperatures to those adapted for low temperatures, occurred between two consecutive evolutionary intermediates. Using a combination of sequence comparisons between ancestral proteins and site-directed mutagenesis analyses, three key amino acid substitutions were identified that enhance low-temperature catalytic activity. Intriguingly, amino acid substitutions that had the most significant impact on activity at low temperatures displayed no discernable effect on thermostability. However, these substitutions markedly reduced the activation energy for catalysis, thereby improving low-temperature activity. Our findings exemplify how ancestral sequence reconstruction can identify residues crucial for adaptation to low temperatures.

## Introduction

Enzymes are the natural proteinaceous catalysts of living organisms. The physical properties of natural enzymes have adapted to the host organism’s habitat and also the competing selection pressures exerted by the cellular environment over a long evolutionary process (1, 2). For instance, enzymes from thermophilic organisms, which thrive in high-temperature environments, tend to display remarkable thermostability and high catalytic activity at elevated temperatures, but their enzymatic activity at low temperatures is often significantly diminished (3–5). In contrast, enzymes isolated from mesophilic organisms tend to be more thermolabile but catalytically more active at low temperatures relative to their thermophilic homologs (6–9). A number of molecular mechanisms that influence the thermostability of an enzyme have been identified by comparative structural analyses between thermophilic and mesophilic enzymes. These studies suggest that the high thermostability of thermophilic enzymes is linked to several factors; (i) increased hydrophobicity and improved packing in the hydrophobic core (10), (ii) increased numbers of intra- or inter-molecular ion-pairs and ion-pair networks found in the native structures (11–13), as well as in the denatured states (14), and (iii) increased flexibility of native structures that results in entropic advantages (15). Comparison of thermophilic and mesophilic or psychrophilic enzymes indicates that the magnitude of their low-temperature activity is often associated with protein surface rigidity (5, 16, 17). However, the precise relationship between surface flexibility and amino acid sequence is poorly understood. Therefore, revealing the mechanism underlying how an enzyme adapts to low temperatures is highly desirable.

Ancestral sequence reconstruction (ASR) is a method of inferring the sequences of ancient genes and proteins that may have been present in extinct organisms based on the phylogenetic analysis of those found in extant genomes (18–20). When the concept of ASR was first proposed by Pauling and Zuckerkandl more than 50 years ago (21) reliable reconstruction of ancestral sequences was unavailable due to the undeveloped nature of the technology at that time. However, the concept was later embodied by advances in phylogenetic analysis technology and computer programs for ancestral sequence inference. The recent colossal expansion of deposited protein sequences in public databases is an essential resource for conducting ASR. The affordability of artificial DNA synthesis in recent years also contributes to advancements in the practical applications of ASR. In particular, inferring and reconstructing amino acid sequences at the root of molecular phylogenetic trees is not only useful in estimating the protein characteristics of ancient organisms (22, 23) but is also invaluable for the synthesis of extremely thermostable enzymes (24, 25). Indeed, most reconstructed ancestral proteins at the deep nodes of phylogenetic trees display high levels of thermostability (24, 25). Given that the most ancestral organism was a thermophile or hyperthermophile that possessed extremely thermostable enzymes (22, 26), extant mesophilic and psychrophilic organisms are thought to have adapted to lower temperatures through the evolutionary process as the Earth’s surface temperature decreased (27).

3-Isopropylmalate dehydrogenase (IPMDH) is involved in the leucine biosynthesis pathway. This enzyme catalyzes the oxidative decarboxylation of D-3-isopropylmalate (D-3-IPM), producing 2-oxoisocaproate, using nicotinamide adenine dinucleotide (NAD^+^) as the electron acceptor. IPMDHs have been the subject of detailed study in terms of their thermostability and low-temperature catalytic activity (28–31). In particular, the relationship between the magnitude of its low-temperature activity and binding affinity to NAD has been investigated (32). Moreover, the principles for improving low-temperature catalytic activity identified from studies of IPMDH have been successfully applied to other dehydrogenases utilizing NAD (or NADP) as a cofactor. As such, the molecular mechanisms of temperature adaptation revealed from the study of IPMDHs are expected to have broad applicability.

Previously, we reconstructed two ancestral IPMDH sequences of the bacterial common ancestor by ASR based on a phylogenetic analysis of extant homologous amino acid sequences (33). These reconstructed ancestral enzymes exhibit higher thermal stability and lower activation energy for catalytic reactions compared to *Thermus thermophilus* IPMDH, which is one of the most intensively studied thermophilic IPMDHs. However, compared to a natural mesophilic IPMDH from *Escherichia coli* (EcIPMDH), these ancestral IPMDHs show reduced catalytic activity at low temperatures. In this study, we explored the evolutionary origins of the enhanced low-temperature catalytic activity found in extant mesophilic enzymes. Specifically, we reconstructed all intermediate sequences along the evolutionary trajectory that connects the oldest ancestor with EcIPMDH. The choice of the trajectory was motivated by the fact that EcIPMDH exhibits a notably higher thermal stability and greater low-temperature activity compared to other mesophilic IPMDHs (31). We found three mutations that contributed to the reduction of activation energy for catalysis that led to a marked improvement in low-temperature IPMDH activity. Thus, this work exemplifies how ASR can be used to identify key mutations that change the catalytic properties of the corresponding enzyme to facilitate adaption to low reaction temperatures.

## Results

### Reconstruction of ancestral IPMDHs

Previously, a multiple sequence alignment of 594 IPMDHs and evolutionarily related proteins from extant species was generated, which was then used to build a maximum likelihood tree (33). A portion of the tree is shown in Fig. S1 (refer to reference 33 for the complete tree) and is schematically represented in Fig. 1A. In this study, the ancestral IPMDH amino acid sequences at each node on the evolutionary trajectory from the most ancient ancestor to EcIPMDH were inferred by using CodeML in PAML (34) and GASP (35) and named Anc01– Anc11. The inferred sequences are given in Data S1 and were aligned with the sequence of EcIPMDH (Fig. S2). Comparison of the amino acid sequences of the most ancient ancestor Anc01 with that of EcIPMDH showed 63% of 353 aligned residues to be identical. Genes encoding the ancestral amino acid sequences were artificially synthesized and engineered for expression in *E. coli* using a pET-based system (Novagen). The recombinant ancestral enzymes were subsequently purified by successive chromatographic steps using HiTrap-Butyl and Superdex 200 Increase columns (Cytiva). As with many other natural IPMDHs, all the ancestral IPMDHs appeared to exist as dimers based on their elution volumes from the gel filtration column during purification.

**Fig. 1.**
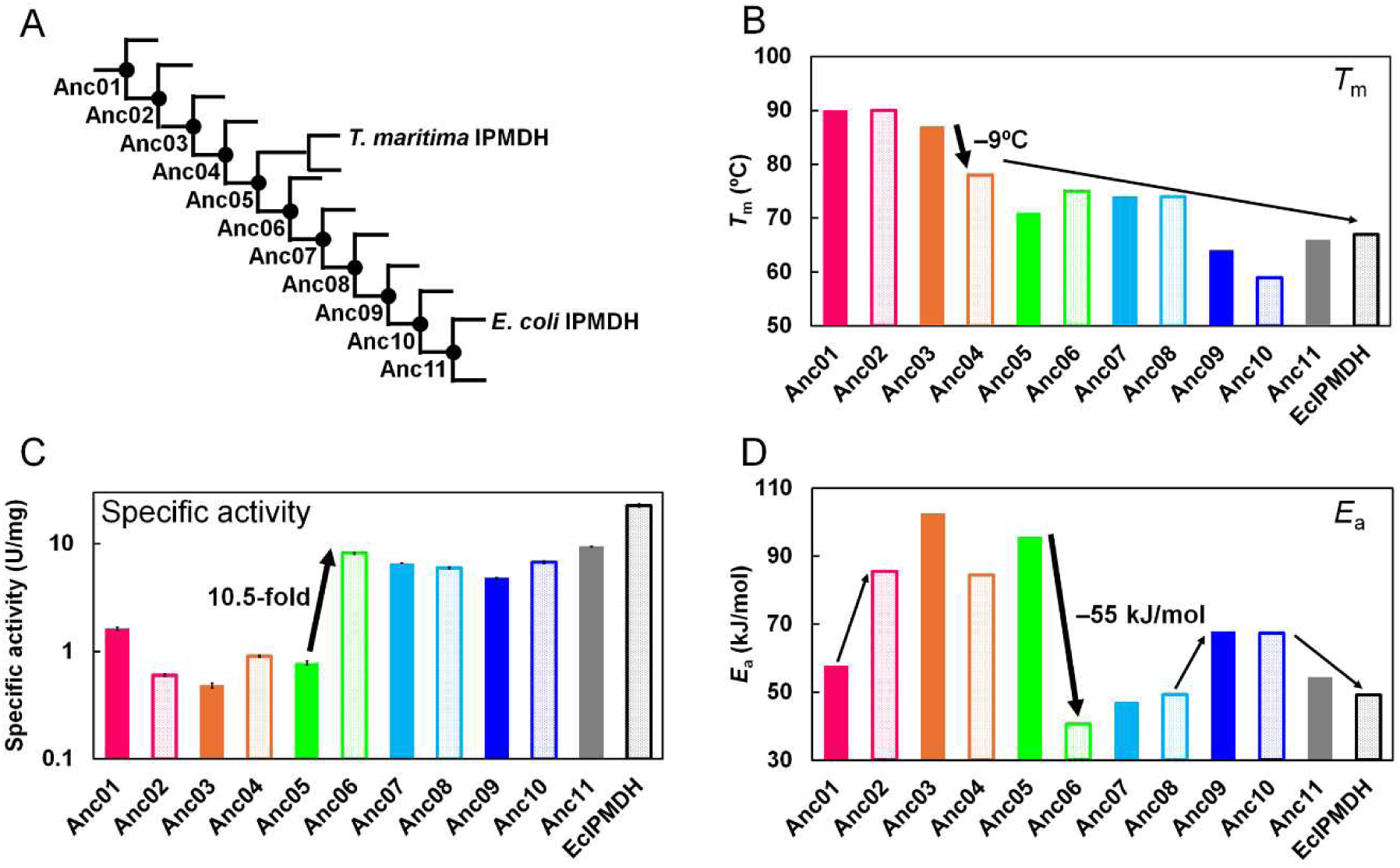
Characterization of ancestral IPMDHs reconstructed in this study. (A) Schematic representation of the trajectory within the phylogenetic tree of IPMDH connecting the oldest ancestor (Anc01) with *E. coli* IPMDH via ten intermediate nodes (Anc02–Anc11). Anc05 is the divergence point between the hyperthermophilic *T. maritima* and the mesophilic *E. coli* IPMDHs. (B) *T*_m_s of the ancestral and *E. coli* IPMDHs. The *T*_m_ values were estimated from the data shown in Fig. S3. The numerical *T*_m_ values are shown in Table S1. (C) Specific activities of the ancestral and *E. coli* IPMDHs at 25°C. The vertical axis uses a logarithmic scale. Each value is the average of three measurements. The error bars represent the standard errors. (D) *E*_a_s of the ancestral and *E. coli* IPMDHs. The *E*_a_ values were estimated from the slopes of the Arrhenius plots shown in Fig. S4B. The numerical *E*_a_ values are given in Table S1.

### Thermal stability

The thermal stability of the ancestral IPMDHs was assessed by tracking ellipticity changes at 222 nm using a 5 μM protein solution. The temperature-induced unfolding curves were normalized by assuming a linear temperature dependence between the baselines of the completely native and fully unfolded states, as depicted in Fig. S3. The unfolding curve for EcIPMDH (31) is also shown in Fig. S3. The unfolding mid-point temperature (*T*_m_) was used to evaluate and compare the thermostability of the IPMDHs (Fig. 1B). As with many reconstructed ancestral proteins, IPMDHs close to the root of the phylogenetic tree exhibited high levels of thermostability (i.e. Anc01, Anc02, Anc03; *T*_m_ ≥ 87°C). Although the *T*_m_ values fluctuated, there was a general decrease along the evolutionary trajectory starting from Anc01 toward the extant mesophilic IPMDH from *E. coli*. The greatest decrease in *T*_m_ was observed between Anc03 (*T*_m_ = 87°C) and Anc04 (*T*_m_ = 78°C).

### Specific activity

The enzymatic activity of the ancestral IPMDHs was evaluated at temperatures ranging from 25–90°C by examining the oxidative decarboxylation of D-3-IPM to yield 2-oxoisocaproate with NAD^+^ serving as the electron acceptor (Data S2). The temperature dependence of the specific activities of the ancestral IPMDHs shows that Anc01, Anc02, and Anc03 exhibited increased specific activities as the reaction temperature arose, reaching their maximum specific activities at 90°C (Fig. S4A). In contrast, the other ancestral IPMDHs and EcIPMDH showed maximum activity at 60–80°C. Typically, cold-adapted enzymes retained approximately 10– 20% of their peak activity at low temperatures, with thermal inactivation occurring before thermal unfolding (36). The mesophilic EcIPMDH functioned optimally at 60°C under the reaction conditions employed and maintained 13% of the maximum activity at 25°C. By contrast, the activities of Anc01, Anc02 and Anc03 at 25°C were reduced to 5.7%, 1.7% and 0.55% of their maximum activities, respectively. Figure 1C compares the specific activities of ancestral IPMDHs and EcIPMDH at 25°C. The largest change was observed between Anc05 and Anc06, where the enzymatic activity increased by 10.5-fold. The relative activity at 25°C compared to the maximum activity was 1.3% for Anc05, which also increased significantly to 28% for Anc06. Although the specific activities at 25°C gradually decreased between Anc06 and Anc09, it increased again from Anc09 toward EcIPMDH.

The activation energies (*E*_a_s) for the specific activities of the ancestral IPMDHs and EcIPMDH were calculated using the slope of the Arrhenius plot (Fig. S4B). A smaller *E*_a_ suggests reduced temperature dependency in catalytic activity, potentially enhancing the catalytic efficiency of cold-adapted enzymes at lower temperatures compared to their counterparts adapted to higher temperatures. Figure 1D shows that Anc01 exhibited a moderate *E*_a_ value (58 kJ/mol), akin to the previously reconstructed ancestral IPMDHs at the node corresponding to the last common ancestor of major bacterial IPMDHs (33). From Anc02 to Anc05, a high *E*_a_ value of greater than 80 kJ/mol was observed but this value decreased sharply between Anc05 and Anc06. Subsequently, the *E*_a_ value increased again up to Anc09, and then gradually declined between Anc10 and EcIPMDH.

### Kinetic parameters at 25°C

The kinetic parameters of catalytic activity for ancestral IPMDHs and EcIPMDH were estimated from steady-state experimental data obtained at 25°C (Fig. 2; Table S1). The *k*_cat_ values followed a trend similar to the specific activities at the same temperature. Notably, akin to the specific activities at 25°C, the *k*_cat_ value increased by 12-fold between Anc05 (0.59 s^-1^) and Anc06 (7.3 s^-1^).

**Fig. 2.**
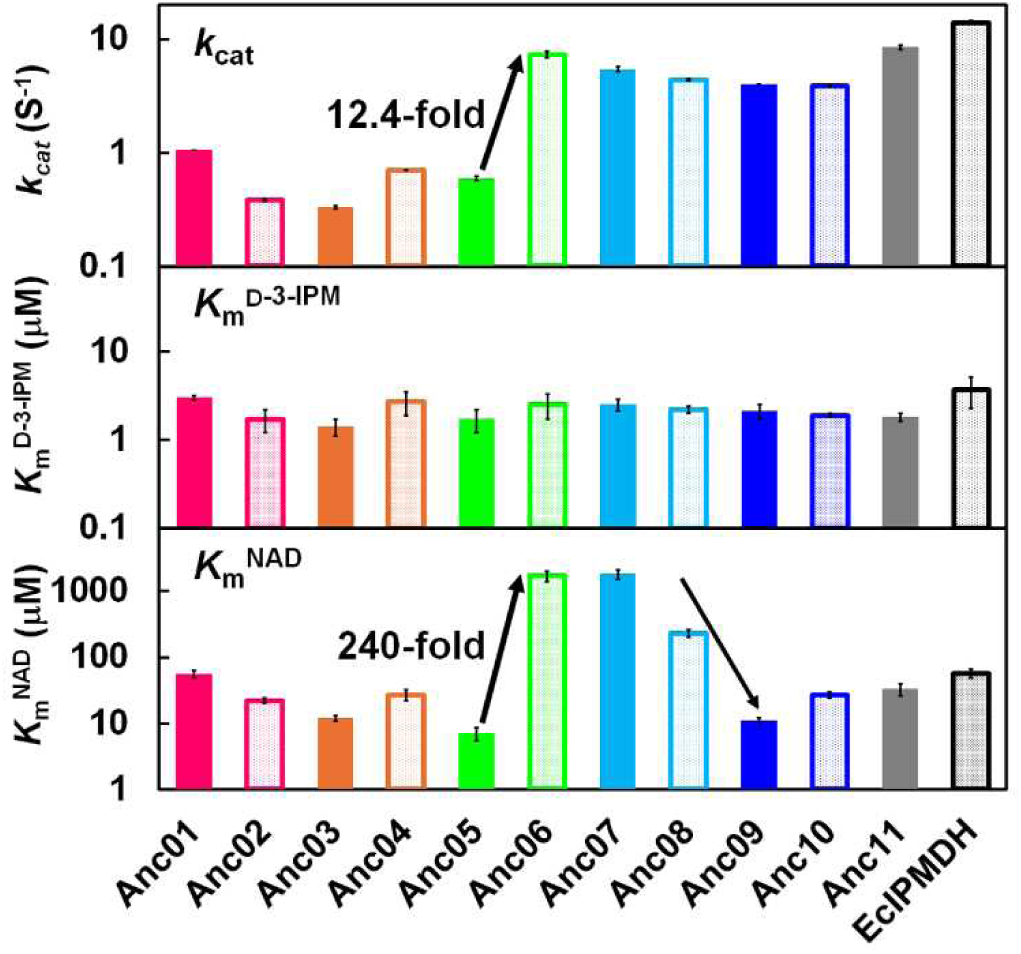
Kinetic parameters of ancestral IPMDHs and EcIPMDH at 25°C. *k*_cat_, *K*_m_^D-3-IPM^, *K*_m_^NAD^, and standard errors were calculated by nonlinear least-square fitting of the steady-state kinetic data to the Michaelis-Menten equation using the Enzyme Kinetics module of SigmaPlot Ver. 14.5 (Systat Software). The vertical axes are shown with a logarithmic scale. The numerical values are given in Table S1.

The *K*_m_^D-3-IPM^ values at 25°C exhibited only slight differences among the ancestral IPMDHs. Conversely, the *K*_m_^NAD^ values displayed significant differences among the enzymes. Specifically, the *K*_m_^NAD^ values sharply increased by 240-fold between Anc05 (7.0 μM) and Anc06 (1700 μM). Therefore, the increase in specific activity and turnover between Anc05 and Anc06 was achieved at the expense of a significant increase in *K*_m_^NAD^. However, a substantial improvement in *K*_m_^NAD^ was observed as evolution progressed from Anc07 (1800 μM) to Anc08 (230 μM), and further to Anc09 (11 μM).

### Exploring key mutations contributing to low-temperature adaptation

Substantial changes in catalytic properties occurred between Anc05 and Anc06. To identify amino acid substitutions that drive the changes in catalytic properties, which make the enzymes better adapted to low reaction temperatures, Anc05 was subjected to site-directed mutagenesis analyses. Initially, amino acid sequences of the ancestral and *E. coli* IPMDHs were compared (Fig. 3A and Fig. S2). This comparison identified five positions (111, 131, 159, 190, and 242) where amino acids were conserved during the evolution from Anc01 to Anc05, replaced between Anc05 and Anc06, and then conserved again through the evolution from Anc06 to EcIPMDH. Additionally, we focused on position 270, where glycine was conserved during the evolution from Anc01 to Anc05, replaced by alanine in Anc06, and then by asparagine in Anc09. Then, we examined how the specific activity at 25°C changed by substituting the amino acids at positions 111, 131, 159, 190, 242, and 270 in Anc05 with those found in the corresponding positions in Anc06. The results showed that the Val112→Ile mutation increased specific activity at 25°C by 3.5-fold, the Val131→Phe mutation by 6.4-fold, the Val159→Phe mutation by 1.4-fold, and the Gly270→Ala mutation by 2.3-fold (Fig. 3B). In contrast, the Val190→Ile and Thr242→Cys mutations did not enhance activity at 25°C.

**Fig. 3.**
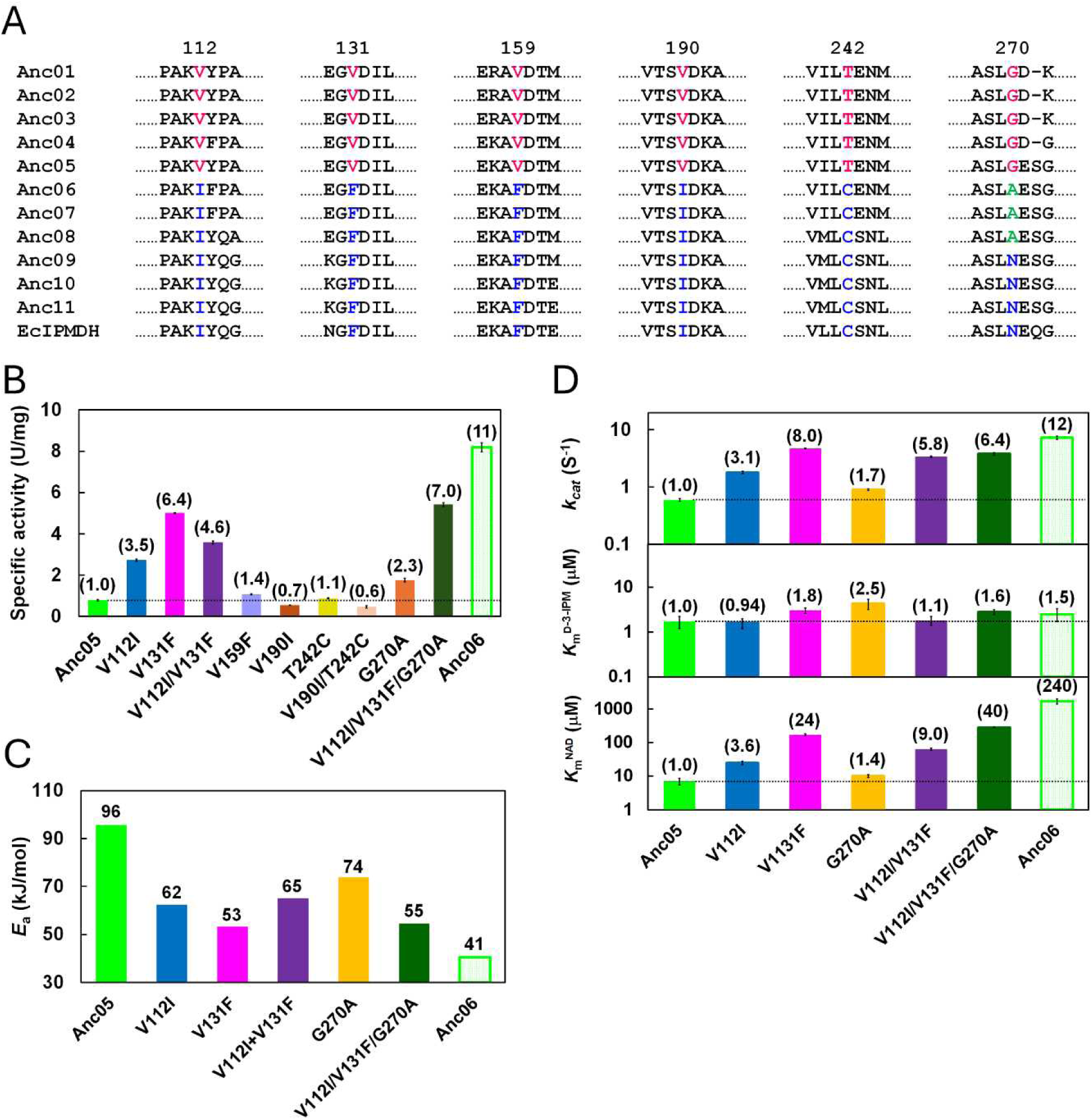
Characterization of the mutants of Anc05. (A) Six mutations that occurred between Anc05 and Anc06. (B) Specific activities of Anc05, its mutants, and Anc06 at 25°C. Values relative to that of Anc05 are given in parentheses above the bars. (C) *E*_a_s of Anc05, its mutants, and Anc06. Numerical values are shown above the bars. The *E*_a_ values were estimated from the slopes of the Arrhenius plots given in Fig. S5B. (D) Kinetic parameters of Anc05, its mutants, and Anc06 at 25°C. *k*_cat_, *K*_m_^D-3-IPM^, *K*_m_^NAD^, and standard errors were calculated by nonlinear least-square fitting of the steady-state kinetic data to the Michaelis-Menten equation using the Enzyme Kinetics module of SigmaPlot Ver. 14.5 (Systat Software). The vertical axes are shown with a logarithmic scale. Values relative to that of Anc05 are shown in parentheses above the bars. The numerical values are given in Table S1.

The predicted tertiary structure of Anc05 using AlphaFold2 (37) indicated that Val112 and Val131 are in close proximity, as are Val190 and Thr242 (Fig. S5). Consequently, we also examined the specific activity at 25°C of the two double mutants V112I/V131F and V190F/T242C. The V112I/V131F mutant exhibited specific activity intermediate between the V112I and V131F single mutants, while the V190F/T242C mutant showed lower activity than the V190F single mutant (Fig. 3B). These results indicated that the simultaneous amino acid substitutions at adjacent sites do not exhibit synergistic effects. We then assessed the effect of the combination of the three beneficial amino acid substitutions (V112I, V131F, G270A). The resulting combined mutant (V112I/V131F/G270A) was slightly more active than the V131F mutant at 25°C, exhibiting 7.0-fold enhanced specific activity relative to Anc05 (Fig. 3B). Based on these findings, we concluded that the three amino acid substitutions were associated, at least in part, with the elevated catalytic activity of Anc06 relative to Anc05 at 25°C.

Next, we investigated the temperature dependence of specific activity for the V112I, V131F, and G270A mutants as well as for the V112I/V131F double mutant and the V112I/V131F/G270A triple mutant (Fig. S6). For three mutants (V112I, V131F, V112I/V131F/G270A) the optimum temperature of catalysis was lower by comparison to that of Anc05. In addition, all mutants exhibited levels of activity between those observed for Anc05 and Anc06 at temperatures ranging from 25 to 40°C.

We also calculated *E*_a_ for the specific activity of each mutant enzyme (Fig. 3C). The Anc05 mutants had reduced *E*_a_ values relative to Anc05. Therefore, the smaller *E*_a_ values are likely responsible for the enhanced catalytic activity of the Anc05 mutants at low temperatures.

In addition, we conducted steady-state kinetic analyses of the Anc05 mutants at 25°C. All mutants showed *k*_cat_ values greater than that of Anc05 (Fig. 3D and Table S1). In particular, the *k*_cat_ value for mutant V131F was 8.0-fold greater relative to Anc05. However, by comparison to Anc05, all mutants showed unfavorable *K*_m_^NAD^ values. Thus, the Val131→Phe mutation improved *k*_cat_ at 25°C because of a trade-off with *K*_m_^NAD^.

### Introduction of the Val→Phe substitution into older and more thermostable ancestors

Our findings from the previous section showed that the Val131→Phe substitution in Anc05 enhanced its specific activity by 6.4-fold at 25°C. Here, we tested whether a corresponding amino acid substitution also improves the low-temperature catalytic activity of older and more thermostable ancestor enzymes. Specifically, we introduced the Val→Phe substitution into Anc01 and Anc03. Anc01 was the oldest and most thermostable ancestor reconstructed in this study and its specific activity was 2.1-fold higher than that of Anc05 (Fig. 1C). Notably, Anc01 had the smaller *E*_a_ value than Anc05 by 31 kJ/mol (Fig. 1D). Anc03 was also more thermostable than Anc05 (Fig. 1B) and had the greatest *E*_a_ value (Fig. 1D) and the smallest specific activity at 25°C (Fig. 1C) among the ancestral IPMDHs. The specific activities of the resulting mutants (Anc01-V128F, Anc03-V128F) as a function of temperature were measured and plotted as shown in Fig. S7. While the Val128*→*Phe mutation did not improve the specific activity of Anc01, Anc03-V128F was more catalytically active relative to Anc03 at temperatures ranging from 25 to 50°C. At 25°C, Anc03-V128F showed enhanced specific activity relative to Anc03 by 5.7-fold (Fig. 4A). The calculated *E*_a_ value of Anc03-V128F was 61 kJ/mol, which was 42 kJ/mol lower than that of Anc03 (Fig. 4B). Thus, Anc03-V128F had improved low-temperature activity due to lowered *E*_a_, akin to that of the corresponding mutant of Anc05.

**Fig. 4.**
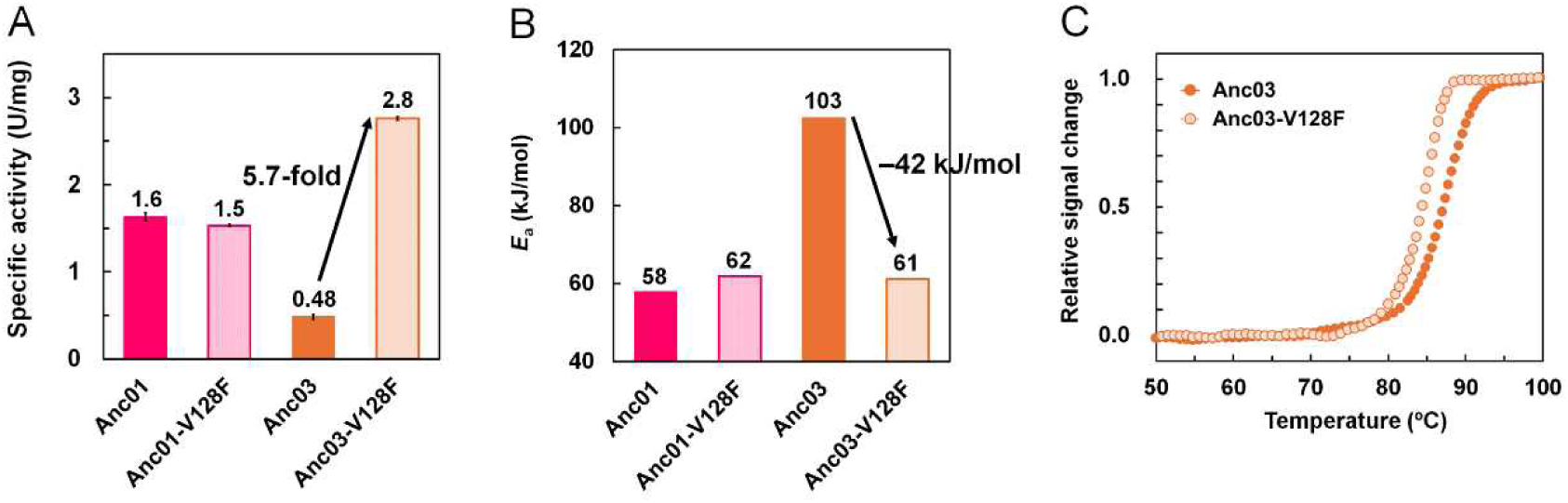
Characterization of the V128F mutants of Anc01 and Anc03. (A) Specific activities of Anc01, Anc03, and their mutants at 25°C. Numerical values are shown above the bars. (B) *E*_a_s of Anc01, Anc03, and their mutants. Numerical values are shown above the bars. The *E*_a_ values were estimated from the slopes of the Arrhenius plots given in Fig. S6B. (C) Temperature-induced unfolding curves of Anc03 and its V128F mutant. The change in ellipticity at 222 nm was monitored as a function of temperature. The temperature was increased at a rate of 1.0°C/min. The samples comprised 5.0 μM protein in 20 mM potassium phosphate (pH 7.6), 0.5 mM EDTA. Each experiment was performed in duplicate with identical unfolding curves within experimental error. The plots were normalized with respect to the baseline of the native and denatured states.

We also performed temperature-induced unfolding experiments for Anc03-V128F. The unfolding curve of Anc03-V128F is shown in Fig. 4C along with that of Anc03 for comparison. The *T*_m_ values of Anc03 and Anc03-V128F were 87°C and 85°C, respectively. Therefore, Anc03-V128F is still a thermostable enzyme although its unfolding temperature is slightly lower than that of Anc03. The results demonstrate that the Val*→*Phe substitution improves the low-temperature activity of Anc03 without significantly compromising its thermostability.

## Discussion

ASR is a way to resurrect plausible ancient sequences from extinct species (18, 19). To date, ASR has been used to test the inevitability of evolutionary pathways of proteins (38), as well as to identify mutations that gave rise to allosteric regulation during evolution (39, 40). ASR is also used in protein engineering to modulate the thermostability and catalytic activity of enzymes (33, 41, 42). In this study, we used ASR to reveal the evolutionary patterns of the thermostability and catalytic activity of IPMDH. Specifically, we found that a limited number of amino acid substitutions, which occurred between two successive intermediate ancestors, contributed to the adaptation of the enzyme to low temperatures. Herschlag and coworkers proposed that temperature adaptation in enzymes is primarily driven by a single mutation of an active-site residue (43). However, amino acid substitutions identified in this study that had an impact on temperature adaptation occurred at positions outside the active site. Wilding and coworkers suggested that ASR can efficiently identify amino acid substitutions remote from the active site that influence thermostability and catalytic activity (44, 45). Thus, ASR may be a way to identify key amino acid residues that are responsible for the adaption of an enzyme to low temperatures.

Most of the reconstructed proteins at the deepest nodes in phylogenetic trees were as thermostable as extant proteins from thermophiles or hyperthermophiles, supporting the idea that the early organisms possessed thermostable proteins and, therefore, that they were thermophilic or hyperthermophilic (22, 46). In addition, other studies expanded the timescale and included more ancestral proteins for reconstruction, suggesting that the oldest ancestor was thermophilic and extant mesophilic and psychrophilic organisms adapted later to lower temperatures during the Precambrian era (26, 27, 47). Indeed, this study revealed the oldest ancestor (Anc01) to have the highest denaturation temperature (*T*_m_=90°C) among the reconstructed ancestral enzymes. However, denaturation temperatures did not decrease smoothly over the course of evolution. Rather, from Anc01 to Anc03, *T*_m_ remained high, followed by a significant decline of 9°C between Anc03 and Anc04. Further along the evolutionary trajectory towards the present time, denaturation temperatures decreased more gradually with some fluctuations. Similar observations have been reported for the evolution of bacterial ribonuclease H1 (RNH). Marqusee and coworkers performed ASR of bacterial RNH to analyze how the thermophilic and mesophilic RNHs evolved from their common ancestor (48). The data showed that the common ancestor had intermediate thermostability between those of the thermophilic and mesophilic RNHs. Intriguingly, the lineage from the common ancestor to the mesophilic descendant of RNH underwent a rapid decrease in thermostability early in the evolutionary process but has subsequently undergone relatively little change.

Akin to changes in *T*_m_, the magnitude of catalytic activity at 25°C did not exhibit a linear change during the evolution from the oldest ancestor to EcIPMDH. Notably, a significant increase in catalytic activity was observed between Anc05 and Anc06, where the specific activity and *k*_cat_ of Anc06 at 25°C were >10-fold greater than those of Anc05. The improvement in *k*_cat_ was accompanied by a 250-fold increase in *K*_m_^NAD^ and a 55 kJ/mol reduction in *E*_a_ for the catalytic reaction. Generally, cold active enzymes show higher specific activity and catalytic turnover, unfavorable *K*_m_, and reduced *E*_a_ relative to their thermophilic counterparts (49–51). Therefore, a substantial change in physical properties involved in the low-temperature adaptation occurred between Anc05 and Anc06, although the largest change in thermostability was observed between Anc03 and Anc04. It is important to note that Anc05 represents the evolutionary divergence point between hyperthermophilic *Thermotoga maritima* and mesophilic *E. coli* IPMDHs (Fig. 1A, Fig S1).

Next, we explored the structural basis underlying the amino acid substitutions responsible for the low-temperature adaptation of IPMDH. AlphaFold2 (37) was used to predict the three-dimensional structure of Anc05, which was then compared with the crystal structure of EcIPMDH (PDB code; 1cm7). Anc05 and EcIPMDH share 68% amino acid sequence identity. The predicted Anc05 structure displayed high concordance with the experimentally determined structure of EcIPMDH (Fig. S8). Superposition of the predicted Anc05 structure, comprising 360 residues, with the structure of EcIPMDH (PDB ID: 1cm7) yielded a C_α_ root-mean-square deviation of 1.92 Å for 342 aligned residues as calculated using PDBeFold (https://www.ebi.ac.uk/msd-srv/ssm/). Most natural IPMDHs are dimeric and their protomers can be divided into two structural domains; domain 1 includes the N- and C-termini, and domain 2 is involved in subunit interaction (Fig. S5). It has been reported that the thermophilic IPMDH from *T. thermophilus* undergoes domain rearrangement upon substrate and cofactor binding (52). The thermophilic IPMDH exhibits reduced activity at room temperature compared to EcIPMDH. This reduced activity in the thermophilic enzyme at room temperature was attributed to more rigid conformational fluctuations, which were necessary for catalysis (29). However, at elevated temperatures, the domain fluctuations increased, thereby enhancing catalytic activity (53). In the predicted structure of Anc05, Val112 and Gly270 are located at an edge of the β-strands that connects the two domains (Fig. 3). In addition, Val131 is spatially close to Val112 in the three-dimensional structure and the side chain of Val131 interacts with that of Val112. Based on these considerations, we anticipate that substitution of Val131 and Val112 with amino acids having larger nonpolar side chains might disrupt the side-chain packing, thereby increasing the fluctuations in domain motion even at lower temperatures. For *E. coli* isocitrate dehydrogenase, which has a similar tertiary structure to IPMDH and catalyze a similar oxidative decarboxylation, the release of a reaction product NADPH was identified as the rate-limiting catalytic step (54). Therefore, the rate-limiting step for the reaction catalyzed by IPMDH is also likely to involve the release of NADH. Because the *K*_m_^NAD+^ values for the Anc05 mutants (Anc05-V131F, Anc05-V112I/V131F, Anc05-V112I/V131F/G270A) are greater than that of the original Anc05 (Fig. 3D) and the affinity of NAD^+^ likely correlates with that of NADH, the amino acid substitutions might increase the fluctuations in domain motion, weakening the IPMDH-NADH binding and thus facilitating liberation of the coenzyme product at low temperatures.

Previously, we improved the low-temperature catalytic activity of *T. thermophilus* IPMDH through amino acid substitutions near the NAD binding site (31, 32). Many of these substitutions weakened the binding of NAD or its reaction product, NADH, by modulating the coenzyme binding site. In contrast, the three mutations identified in this study are distant from the active site. This differs from the previous studies that required mutations around the NAD binding site. Therefore, the present work identified an alternative strategy for promoting low-temperature adaptation in IPMDH. Specifically, we propose that, for enzymes whose magnitude of fluctuations in domain motion influences the magnitude of catalytic activity, the base positions of the segments connecting the domains serve as potential targets for mutation in order to enhance catalytic activity at low temperatures.

While the physical and catalytic properties vary in naturally occurring enzymes, the trade-off between thermostability and low-temperature activity are disputed (5, 50). Among the ancestral and *E. coli* IPMDHs, an inverse-correlation between their logarithms of specific activities at 25°C and *T*_m_s were observed (Fig. 5A and Table 1). Therefore, enhancement of the low-temperature activity was achieved, at least in part, at the expense of thermostability in the course of IPMDH evolution. The inverse correlation was evident up to 60°C, but at 70°C, only a weak correlation was observed. These results imply that the magnitude of catalytic activity is inversely related to thermostability, particularly at temperatures where thermal denaturation does not occur. Notably, correlation coefficient (*R*) values indicate that the variability of the data at 25°C is greatest among those between 25°C and 60°C, suggesting that as reaction temperature decreases, factors independent of thermostability play a more prominent role in determining the magnitude of enzymatic activity. Interestingly, a slight improvement in thermostability was observed between Anc05 and Anc06, coinciding with the most pronounced increase in low-temperature activity. Accordingly, as suggested previously (55, 56), it is apparent that the catalytic activity of an enzyme at lower temperatures can be improved without necessarily compromising thermostability.

**Fig. 5.**
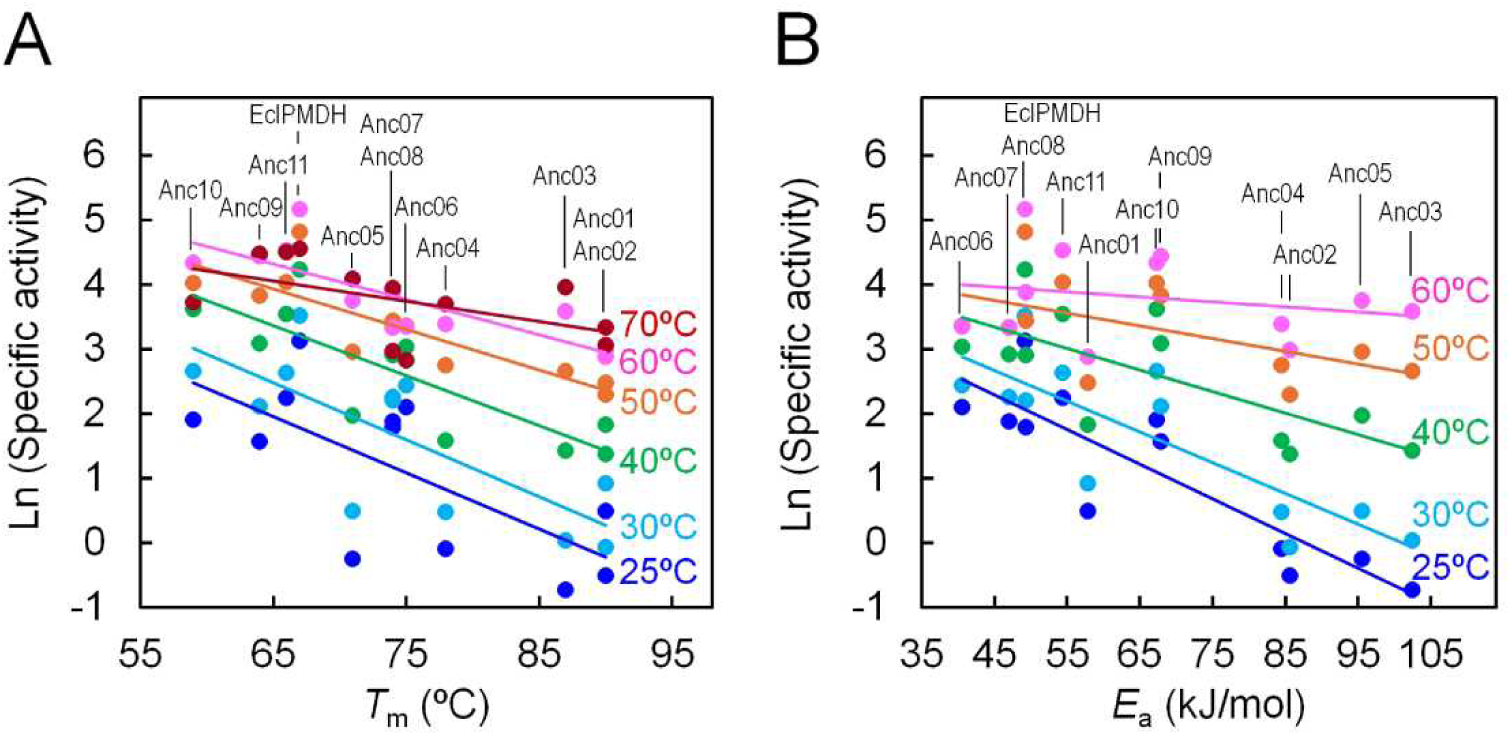
The specific activities of ancestral and *E. coli* IPMDHs at various temperatures shown with a logarithmic scale as a function of their *T*_m_s (A) or *E*_a_s (B). The linear approximation of the plot is given as a solid line. Correlation coefficients and slopes are listed in Table 1.

**Table 1.** Relationship between *T*_m_ or *E*_a_ and the specific activities at various temperatures on a logarithmic scale of ancestral and *E. coli* IPMDHs. Correlation coefficients (*R*) and slopes were obtained from the linear approximation of the plots shown in Fig. 5.

| Temperature<br>(°C) | $T_m$ vs. Specific activity | | $E_a$ vs. Specific activity | |
| --- | --- | --- | --- | --- |
| | $R$ | Slope | $R$ | Slope |
| 25 | −0.69 | −0.087 | −0.87 | −0.054 |
| 30 | −0.75 | −0.089 | −0.82 | −0.048 |
| 40 | −0.82 | −0.077 | −0.72 | −0.033 |
| 50 | −0.85 | −0.063 | −0.55 | −0.020 |
| 60 | −0.81 | −0.055 | −0.23 | −0.008 |
| 70 | −0.53 | −0.031 | — | — |

The logarithms of specific activities at 25°C correlate more strongly with the *E*_a_s (Figs. 5B and Table 1). Given that *E*_a_ represents the temperature dependence of catalytic activity, it is unsurprising that the inverse correlation strengthens as the reaction temperature decreases. However, Anc01 exhibited relatively low *E*_a_ yet also displayed a low specific activity. Similarly, EcIPMDH deviated significantly above the linear approximation at 25°C, as well as at other temperatures. This observation suggests that a lower *E*_a_ alone cannot account for the magnitude of low-temperature activity. Nevertheless, reducing *E*_a_ remains a key strategy in enzyme engineering aimed at enhancing low-temperature activity. Further investigation is needed to clarify additional mechanisms influencing catalytic activity at low temperatures.

In conclusion, our results demonstrate that ancestral sequence reconstruction is an invaluable methodology for exploring the molecular mechanisms underlying the evolutionary adaptation of an enzyme to low temperatures. Through comprehensive reconstruction of the ancestral sequences along the evolutionary trajectory from the thermophilic ancestor of IPMDHs to the extant mesophilic enzyme, three key mutations involved in the adaption of the enzyme to low temperatures were identified. Notably, this work identified specific interactions that contributed to the smaller *E*_a_ values, which was responsible for enhanced low-temperature activities. Moreover, we revealed the evolutionary patterns of thermostability, temperature dependence of catalytic activity, activation energy for catalysis, and steady-state kinetic parameters. Nonetheless, one outstanding question remains. Why did the key amino acid substitution (Val131*→*Phe) not have the same effect on all ancestral enzymes? Specifically, the Val*→*Phe substitution enhanced the low-temperature activities of Anc03 and Anc05, but the mutation had no effect on that of Anc01. Thus, identical amino acid substitutions may not have the same effect in different enzymes even if they are evolutionarily closely related. Noteworthy, Anc03 and Anc05 exhibited significantly large *E*_a_ values by comparison to Anc01. This observation may be linked to the apparent disparity. A major aim of future studies will be to elucidate a set of universal structure-based rules that will identify common amino acid substitutions for enhancing low-temperature activity in different enzymes.

## Materials and Methods

### Ancestral amino acid sequence inference

Previously, we built a maximum likelihood tree of extant IPMDHs and their evolutionarily related enzymes (33). The phylogenetic tree comprised 594 sequences (435 bacterial and 89 archaeal sequences). In this study, we inferred ancestral amino acid sequences of all nodes along the evolutionary lineage that connects the bacterial common ancestral IPMDH and EcIPMDH. To infer the ancestral amino acid sequences, we used CodeML in PAML (34) with the LG + Gamma (eight-class) model. We also used GASP (35) to estimate the location of gaps in the ancestral sequences. The inferred sequences are shown in Fig. S2

### Cloning and expression

The inferred ancestral amino acid sequences of IPMDH were reverse-translated to generate their gene sequences. The genes were synthesized by Eurofins Genomics. NdeI and BamHI restriction recognition sites were engineered upstream and downstream of the synthesized genes, respectively. The synthetic genes were digested with NdeI and BamHI and then cloned into the NdeI-BamHI sites of plasmid pET23a(+) (Merck). To express the ancestral IPMDHs, the resulting plasmids were used to transform *E. coli* Rosetta2(DE3) and the transformants were cultivated in LB medium supplemented with ampicillin (150 μg/ml). Overexpression was induced using the Overnight Express Autoinduction system I (Merck). After overnight culture at 37°C, cells were harvested by centrifugation and the cell pellets stored at –80°C.

### Enzyme purification

To purify ancestral IPMDHs, cell pellets were suspended in 20 mM Tris-HCl (pH 8.0), 1 mM EDTA and disrupted by sonication. The soluble fractions were isolated after centrifugation at 60,000 × *g* for 20 min. The supernatants were individually heated at 60°C or 70°C for 15 min and centrifuged at 60,000 × *g* for 20 min. The clarified cell-free extracts were then subjected to hydrophobic interaction chromatography using a HiTrapButyl column (GE Healthcare Bioscience) followed by gel filtration chromatography on a Superdex 200 Increase column (GE Healthcare Bioscience). The homogeneities of the purified enzymes were found to be >95% as assessed by sodium dodecyl sulfate-polyacrylamide gel electrophoresis.

### Thermal stability measurements

Protein concentrations were determined using the A_280_ values of protein-containing solutions. Molar extinction coefficient of each enzyme was calculated based on the amino acid sequence as described by Pace and colleagues (57) who followed the procedure of Gill and von Hippel (58).

The thermal denaturation profile for each enzyme was monitored by measuring the change in ellipticity at 222 nm using a J-1100 spectropolarimeter (Jasco) equipped with a programmable temperature controller and a pressure-proof cell compartment to prevent the aqueous solution from bubbling and evaporating at high temperatures. Each enzyme was diluted to a final concentration of 5.0 μM with 20 mM potassium phosphate (pH 7.6), 1 mM EDTA. Measurements were performed using a 0.1 cm pathlength quartz cuvette, and the temperature was increased at a rate of 1.0°C/min.

### Activity measurements

The specific activity of each IPMDH was measured by monitoring the change in absorbance at 340 nm, which corresponded to the rate of production of NADH. The assay buffer consisted of 50 mM HEPES (pH 8.0), 100 mM KCl, 5.0 mM MgCl_2_, 0.40 mM DL-3-isopropylmalate (DL-3-IPM), and 5.0 mM NAD^+^. One enzyme unit corresponded to 1 μmol NADH formed per min.

The values of *K_m_*^NAD+^ and *k_cat_* were determined using steady-state kinetic data in assay buffer where the NAD^+^ concentration was varied between 1.0 and 5000 μM. To determine the *K_m_* ^D-3-IPM^ value, the substrate concentration was varied between 1.0 and 30 μM, while the coenzyme concentration was fixed at 5000 μM. The kinetic constants were obtained by nonlinear least-square fitting of the steady-state velocity data to the Michaelis-Menten equation using the Enzyme Kinetics module of SigmaPlot 14.5 (Systat Software).

### Site-directed mutagenesis

Site-directed mutagenesis of ancestral IPMDH genes was conducted using the splicing-by-overlap-extension PCR method (59). The nucleotide sequences of the synthetic oligonucleotide primers are given in Table S2. To amplify mutagenized genes, the PCR-reaction mixture contained 1× PCR buffer for KOD-plus polymerization (Toyobo), 1.0 mM MgSO_4_, 0.20 mM each of the dNTPs, 0.40 μM each of the synthetic oligonucleotides, and 1.0 unit KOD-plus DNA polymerase. The PCR temperature cycling program was as follows: step 1, 95°C, 3 min; step 2, 95°C, 30 s; step 3, 55°C, 30 s; step 4, 68°C, 1.0 min; steps 2–4 were repeated 25 times. After amplification, the PCR product was purified and then digested with NdeI and BamHI (New England Biolabs), followed by cloning into the NdeI-BamHI sites of pET23a. Expression, purification and analyses of each mutant protein was performed as described above.

## Acknowledgments

This work was supported by JSPS KAKENHI (Grant number 23K17851) to SA.

## References

1. S. Jemli, D. Ayadi-Zouari, H.B. Hlima, S. Bejar, Biocatalysts: application and engineering for industrial purposes. Crit. Rev. Biotechnol. 36, 246–258 (2016).

2. S. Prasad, I. Roy, Converting enzymes into tools of industrial importance. Recent Pat. Biotechnol. 12, 33–56 (2018).

3. C. Vieille, G.J. Zeikus, Hyperthermophilic enzymes: sources, uses, and molecular mechanisms for thermostability. Microbiol. Mol. Biol. Rev. 65,1–43 (2001).

4. P.A. Fields, Review: Protein function at thermal extremes: balancing stability and flexibility. Comp. Biochem. Physiol. A Mol. Integr. Physiol. 129, 417–431 (2001).

5. G. Feller, Protein stability and enzyme activity at extreme biological temperatures. J. Phys. Condens. Matter. 22, 323101 (2010).

6. M. Mangiagalli, S. Brocca, M. Orlando, M. Lotti, The “cold revolution”. Present and future applications of cold-active enzymes and ice-binding proteins. N. Biotechnol. 55, 5–11 (2020).

7. T. Collins, R. Margesin, Psychrophilic lifestyles: mechanisms of adaptation and biotechnological tools. Appl. Microbiol. Biotechnol. 103, 2857–2871 (2019).

8. K.S. Siddiqui, Some like it hot, some like it cold: Temperature dependent biotechnological applications and improvements in extremophilic enzymes. Biotechnol. Adv. 33, 1912–1922 (2015).

9. D. Georlette, et al. Some like it cold: biocatalysis at low temperatures. FEMS Microbiol. Rev. 28, 25–42 (2004).

10. M.M. Gromiha, M.C. Pathak, K. Saraboji, E.A. Ortlund, E.A. Gaucher, Hydrophobic environment is a key factor for the stability of thermophilic proteins. Proteins 81, 715–721 (2013).

11. C. Vetriani, et al. Protein thermostability above 100 degreesC: a key role for ionic interactions. Proc. Natl. Acad. Sci. U.S.A. 95, 12300–12305 (1998).

12. T. Tanaka, et al. Hyper-thermostability of CutA1 protein, with a denaturation temperature of nearly 150 degrees C. FEBS Lett. 580, 4224–4230 (2006).

13. S. Irumagawa, et al. Rational thermostabilisation of four-helix bundle dimeric *de novo* proteins. Sci. Rep. 11, 7526 (2021).

14. K. Yutani, Y. Matsuura, H. Naitow, Y. Joti, Ion-ion interactions in the denatured state contribute to the stabilization of CutA1 proteins. Sci Rep 8, 7613 (2018).

15. Z. Liu, et al. Entropic contribution to enhanced thermal stability in the thermostable P450 CYP119. Proc. Natl. Acad. Sci. U.S.A. 115, E10049–e10058 (2018).

16. G.V. Isaksen, J. Aqvist, B.O. Brandsdal, Protein surface softness is the origin of enzyme cold-adaptation of trypsin. PLoS Comput. Biol. 10, e1003813 (2014).

17. G.V. Isaksen, J. Aqvist, B.O. Brandsdal, Enzyme surface rigidity tunes the temperature dependence of catalytic rates. Proc. Natl. Acad. Sci. U.S.A. 113, 7822–7827 (2016).

18. R. Merkl, R. Sterner, Ancestral protein reconstruction: techniques and applications. Biol. Chem. 397, 1–21 (2016).

19. E.A. Gaucher, J.T. Kratzer, R.N. Randall, Deep phylogeny--how a tree can help characterize early life on Earth. Cold Spring Harb. Perspect. Biol. 2, a002238 (2010).

20. J.W. Thornton, Resurrecting ancient genes: experimental analysis of extinct molecules. Nat. Rev. Genet. 5, 366–375 (2004).

21. Z.E. Pauling L, Chemical paleogenetics: molecular restoration studies of extinct forms of life. Acta Chem. Scand. 17, S9–S16 (1963).

22. S. Akanuma, et al. Experimental evidence for the thermophilicity of ancestral life. Proc. Natl. Acad. Sci. U.S.A. 110, 11067–11072 (2013).

23. F. Busch, et al. Ancestral Tryptophan Synthase Reveals Functional Sophistication of Primordial Enzyme Complexes. Cell Chem. Biol. 23, 709–715 (2016).

24. L.C. Wheeler, S.A. Lim, S. Marqusee, M.J. Harms, The thermostability and specificity of ancient proteins. Curr. Opin. Struct. Biol. 38, 37–43(2016).

25. R.E.S. Thomson, S.E. Carrera-Pacheco, E.M.J. Gillam, Engineering functional thermostable proteins using ancestral sequence reconstruction. J. Biol. Chem. 298, 102435 (2022).

26. V. Nguyen, et al. Evolutionary drivers of thermoadaptation in enzyme catalysis. Science 355, 289–294 (2017).

27. A.K. Garcia, J.W. Schopf, S.I. Yokobori, S. Akanuma, A. Yamagishi, Reconstructed ancestral enzymes suggest long-term cooling of Earth’s photic zone since the Archean. Proc. Natl. Acad. Sci. U.S.A. 114, 4619–4624 (2017).

28. G. Wallon, et al. Crystal structures of *Escherichia coli* and *Salmonella typhimurium* 3-isopropylmalate dehydrogenase and comparison with their thermophilic counterpart from *Thermus thermophilus*. J. Mol. Biol. 266, 1016–1031 (1997).

29. A. Svingor, J. Kardos, I. Hajdú, A. Németh, P. Závodszky, Adjustment of conformational flexibility is a key event in the thermal adaptation of proteins. Proc. Natl. Acad. Sci. U.S.A. 95, 7406–7411 (1998).

30. A. Svingor, J. Kardos, I. Hajdú, A. Németh, P. Závodszky, A better enzyme to cope with cold. Comparative flexibility studies on psychrotrophic, mesophilic, and thermophilic IPMDHs. J. Biol. Chem. 276, 28121–28125 (2001).

31. S. Akanuma, et al. Establishment of mesophilic-like catalytic properties in a thermophilic enzyme without affecting its thermal stability. Sci. Rep. 9, 9346 (2019).

32. S. Hayashi, S. Akanuma, W. Onuki, C. Tokunaga, A. Yamagishi, Substitutions of coenzyme-binding, nonpolar residues improve the low-temperature activity of thermophilic dehydrogenases. Biochemistry 50, 8583–8593 (2011).

33. R. Furukawa, W. Toma, K. Yamazaki, S. Akanuma, Ancestral sequence reconstruction produces thermally stable enzymes with mesophilic enzyme-like catalytic properties. Sci. Rep. 10, 15493 (2020).

34. Z. Yang, PAML 4: phylogenetic analysis by maximum likelihood. Mol. Biol. Evol. 24, 1586–1591 (2007).

35. R.J. Edwards, D.C. Shields, GASP: Gapped Ancestral Sequence Prediction for proteins. BMC Bioinformatics 5, 123 (2004).

36. G. Feller, Psychrophilic enzymes: from folding to function and biotechnology. Scientifica 2013, 512840 (2013).

37. J. Jumper, et al. Highly accurate protein structure prediction with AlphaFold. Nature 596, 583–589 (2021).

38. T.N. Starr, L.K. Picton, J.W. Thornton, Alternative evolutionary histories in the sequence space of an ancient protein. Nature 549, 409–413 (2017).

39. A. Hadzipasic, et al. Ancient origins of allosteric activation in a Ser-Thr kinase. Science 367, 912–917 (2020).

40. M. Schupfner, K. Straub, F. Busch, R. Merkl, R. Sterner, Analysis of allosteric communication in a multienzyme complex by ancestral sequence reconstruction. Proc. Natl. Acad. Sci. U.S.A. 117, 346–354 (2020).

41. S. Nakano, Y. Minamino, F. Hasebe, S. Ito, Deracemization and Stereoinversion to Aromatic D-Amino Acid Derivatives with Ancestral L-Amino Acid Oxidase. ACS Catalysis 9, 10152–10158 (2019).

42. Y. Gumulya, et al. Engineering highly functional thermostable proteins using ancestral sequence reconstruction. Nat. Catal. 1, 878–888 (2018).

43. M.M. Pinney, et al. Parallel molecular mechanisms for enzyme temperature adaptation. Science 371, eaay2784 (2021).

44. M. Wilding, N. Hong, M. Spence, A.M. Buckle, C.J. Jackson, Protein engineering: the potential of remote mutations. Biochem. Soc. Trans. 47, 701–711 (2019).

45. M. Wilding, et al. Reverse engineering: transaminase biocatalyst development using ancestral sequence reconstruction. Green Chem. 19, 5375–5380 (2017).

46. E.A. Gaucher, J.M. Thomson, M.F. Burgan, S.A. Benner, Inferring the palaeoenvironment of ancient bacteria on the basis of resurrected proteins. Nature 425, 285–288 (2003).

47. E.A. Gaucher, S. Govindarajan, O.K. Ganesh, Palaeotemperature trend for Precambrian life inferred from resurrected proteins. Nature 451, 704–707 (2008).

48. K.M. Hart, et al. Thermodynamic system drift in protein evolution. PLoS Biol. 12, e1001994 (2014).

49. S. Bjelic, B.O. Brandsdal, J. Aqvist, Cold adaptation of enzyme reaction rates. Biochemistry 47, 10049–10057 (2008).

50. K.S. Siddiqui, R. Cavicchioli, Cold-adapted enzymes. Annu. Rev. Biochem. 75, 403–433 (2006).

51. A.O. Smalas, H.K. Leiros, V. Os, N.P. Willassen, Cold adapted enzymes. Biotechnol. Annu. Rev. 6, 1–57 (2000).

52. S. Kadono, et al. Ligand-induced changes in the conformation of 3-isopropylmalate dehydrogenase from *Thermus thermophilus*. J. Biochem. 118, 745–752 (1995).

53. I. Hajdú, A. Szilágyi, J. Kardos, P. Závodszky, A link between hinge-bending domain motions and the temperature dependence of catalysis in 3-isopropylmalate dehydrogenase. Biophys. J. 96, 5003–5012 (2009).

54. A.M. Dean, D.E. Koshland, Jr., Kinetic mechanism of *Escherichia coli* isocitrate dehydrogenase. Biochemistry 32, 9302–9309 (1993).

55. Miller, S. R. An appraisal of the enzyme stability-activity trade-off. Evolution 71, 1876–1887 (2017)

56. K.S. Siddiqui, Defying the activity-stability trade-off in enzymes: taking advantage of entropy to enhance activity and thermostability. Crit. Rev. Biotechnol. 37, 309–322 (2017)

57. C.N. Pace, F. Vajdos, L. Fee, G. Grimsley, T. Gray, How to measure and predict the molar absorption coefficient of a protein. Protein Sci. 4, 2411–2423 (1995).

58. S.C. Gill, P.H. von Hippel, Calculation of protein extinction coefficients from amino acid sequence data. Anal. Biochem. 182, 319–326 (1989).

59. R.M. Horton, Z. Cai, S.M. Ho, L.R. Pease, Gene splicing by overlap extension: tailor-made genes using the polymerase chain reaction. BioTechniques 8(5):528–535 (November 1990). Biotechniques 54, 129–133 (2013).

