## Supplementary materials for "Insights into the low-temperature adaptation of an enzyme as studied through ancestral sequence reconstruction"

**This PDF file includes:**

Figures S1 to S8
Tables S1 and S2
Data S1
Legend for Data S2

**Other supplementary materials for this manuscript include the following:**

Data S2

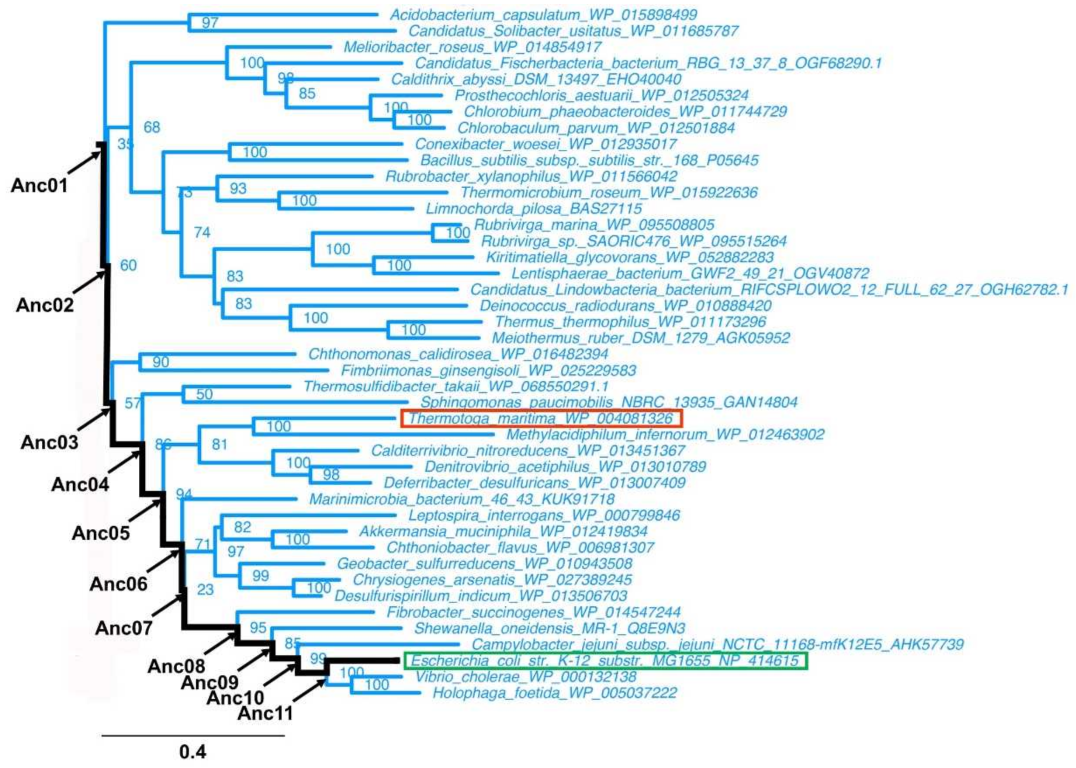


**Figure S1. A portion of the phylogenetic tree of bacterial IPMDHs constructed previously (33).** Organism names and database accession numbers are shown. The arrows mark the nodes corresponding to the positions of the ancestral IPMDH. The hyperthermophilic IPMDH for *Thermotoga maritima* and the mesophilic IPMDH from *Escherichia coli* are boxed. The number on each node shows the RELL bootstrap probability. For the complete tree, see Reference 33. The scale bar represents 0.4 substitutions per site.

```
          1                                                         59
Anc01     M--TYKIAVLPGDGIGPEVVAEAVKVLEAVAEKFGLEFEFEEALIGGAAIDATGTPLPEE
Anc02     M--TYKIAVLPGDGIGPEVVAEAVKVLEAVAEKFGLEFEFEEALIGGAAIDAHGTPLPEE
Anc03     M--TYKIAVLPGDGIGPEVVAEAVKVLEAVAEKFGLEFEFEEALIGGAAIDAHGHPLPEE
Anc04     M--TYKIAVLPGDGIGPEVMAEALKVLEAVSKKFGLEFEFEEALVGGAAIDAHGHPLPEE
Anc05     M-KTYKIAVLPGDGIGPEVMAEALKVLDAVSKKFGLEFEYEEAHVGGAAIDNHGTPLPEE
Anc06     M-KTYKIAVLPGDGIGPEVMAEALKVLDAVSKKFGLTFEYEEAHVGGAAIDNHGSPLPEE
Anc07     M-KTYKIAVLPGDGIGPEVMAEALKVLDAVSKKFGLTFEYEEAHVGGAAIDNHGSPLPEE
Anc08     MTKTYKIAVLPGDGIGPEVMAEALKVLDAVSKKFGLNFEYEEANVGGAAIDNHGSPLPES
Anc09     MTKTYKIAVLPGDGIGPEVMAEALKVLDAVEQKFGLNFEYSEYDVGGAAIDNHGCPLPES
Anc10     MTKTYKIAVLPGDGIGPEVMAEALKVLDAVEQKFGLNFEYSEYDVGGAAIDNHGCPLPES
Anc11     MTKTYKIAVLPGDGIGPEVMAEALKVLDAVEQKFGLRFTYSEYDVGGAAIDNHGCPLPES
EcIPMDH   MSKNYHIAVLPGDGIGPEVMTQALKVLDAVRNRFAMRITTSHYDVGGAAIDNHGQPLPPA

          60                                                  112   119
Anc01     TLEVCKESDAVLLGAVGGPKWDNLPPDLRPERG-LLKLRKALGLFANLRPAKVYPALVDA
Anc02     TLEVCKESDAVLLGAVGGPKWDNLPPDLRPERG-LLKLRKALGLFANLRPAKVYPALVDA
Anc03     TLEVCKESDAVLLGAVGGPKWDNLPPELRPERGALLPLRKALGLFANLRPAKVYPALVDA
Anc04     TLEVCEESDAILFGAVGGPKWENLPPEQQPERGALLPLRKAFGLFANLRPAKVFPALVDA
Anc05     TLKICEESDAILFGSVGGPKWENLPPEQQPERGALLPLRKHFGLFANLRPAKVYPALVDA
Anc06     TLKICEESDAILFGSVGGPKWENLPPEQQPERGALLPLRKHFGLFANLRPAKIFPALAHA
Anc07     TLKICEESDAILFGSVGGPKWENLPPEQQPERGALLPLRKHFGLFANLRPAKIFPALAHA
Anc08     TLKLCEESDAILFGSVGGPKWEHLPPNQQPERGALLPLRKHFKLFCNLRPAKIYQALAHA
Anc09     TLKACEESDAILFGSVGGPKWEHLPPNQQPERGALLPLRKHFKLFCNLRPAKIYQGLEHA
Anc10     TLKACEESDAILFGSVGGPKWEHLPPNQQPERGALLPLRKHFKLFCNLRPAKIYQGLEHA
Anc11     TLKACEEADAVLFGSVGGPKWEHLPPNQQPERGALLPLRKHFKLFCNLRPAKIYQGLEHF
EcIPMDH   TVEGCEQADAVLFGSVGGPKWEHLPPDQQPERGALLPLRKHFKLFSNLRPAKIYQGLEAF

         120        131                          159                177
Anc01     SPLKPEVV-EGVDILVVRELTGGIYFGQPRGID-KG-NERAVDTMVYTRSEIERIARVAF
Anc02     SPLKPEVV-EGVDILVVRELTGGIYFGQPRGRD-EG--ERAVDTMVYTRSEIERIARVAF
Anc03     SPLKPEIV-EGVDILVVRELTGGIYFGQPRGRD-EG--ERAVDTMVYTRSEIERIARVAF
Anc04     SPLKPEIV-EGVDILVVRELTGGIYFGQPKGRD-EG-KEKAVDTMVYSRSEIERIARVAF
Anc05     SPLKEEIVGEGVDILVVRELTGGIYFGQPKGRD-EG-EEKAVDTMVYSRSEIERIARVAF
Anc06     SPLKEEIVGEGFDILVVRELTGGIYFGQPKGREGEGEEEKAFDTMVYSRSEIERIARVAF
Anc07     SPLKEEIVGEGFDILVVRELTGGIYFGQPKGREGEGEEEKAFDTMVYSRSEIERIARVAF
Anc08     SPLRADIVGEGFDILVVRELTGGIYFGQPKGREGEGEEEKAFDTMVYSRSEIERIARIAF
Anc09     SPLRADISAKGFDILCVRELTGGIYFGQPKGREGEGEHEKAFDTMVYSRREIERIARIAF
Anc10     SPLRADISAKGFDILCVRELTGGIYFGQPKGREGEGEHEKAFDTEVYSRREIERIARIAF
Anc11     SPLRADISAKGFDILCVRELTGGIYFGQPKGREGEGEHEKAFDTEVYHRFEIERIARIAF
EcIPMDH   CPLRADIAANGFDILCVRELTGGIYFGQPKGREGSGQYEKAFDTEVYHRFEIERIARIAF

         178          190                                          237
Anc01     ELARKRRKKVTSVDKANVLESSQLWREVVTEVAKEYPDVELEHMYVDNCAMQLVRNPRQF
Anc02     EVARKRRKKVTSVDKANVLETSQLWREVVTEVAKEYPDVELEHMYVDNCAMQLVRDPRQF
Anc03     EVARKRRKKVTSVDKANVLETSQLWREVVTEVAKEYPDVELEHMYVDNCAMQLVRDPRQF
Anc04     EVARKRRKKVTSVDKANVLTTSVLWREVVTEVAKDYPDVELEHMYVDNCAMQLVRDPRQF
Anc05     EVARKRRKKVTSVDKANVLTTSVLWREVVTEVAKDYPDVELEHMYVDNAAMQLVRNPRQF
Anc06     EVARKRRKKVTSIDKANVLTTSVLWREVVTEVAKDYPDVELEHMYVDNAAMQLVRNPRQF
Anc07     EAARKRRKKVTSIDKANVLTTSVLWREVVTEVAKDYPDVELEHMYVDNAAMQLVRNPRQF
Anc08     EAARLRRKKVTSIDKANVLTTSVLWREVVTEVAKDYPDVELEHMYVDNAAMQLVKNPRQF
Anc09     EAARLRRKKVTSIDKANVLASSVLWREVVTEVAKDYPDVELEHMYIDNATMQLVKNPSQF
Anc10     ESARLRRKKVTSIDKANVLASSILWREVVTEVAKDYPDVELEHMYIDNATMQLVKNPSQF
Anc11     ESARLRRKKVTSIDKANVLQSSILWREVVTEVAKDYPDVELSHMYIDNATMQLIKDPSQF
EcIPMDH   ESARKRRHKVTSIDKANVLQSSILWREIVNEIATEYPDVELAHMYIDNATMQLIKDPSQF
```

```
         238 242                         270                        297
Anc01      DVILTENMFGDILSDEAAMLTGSLGMLPSASLGD-KVGLYEPVHGSAPDIAGQGIANPIA
Anc02      DVILTENMFGDILSDEAAMLTGSLGMLPSASLGD-KFGLYEPVHGSAPDIAGQGIANPIA
Anc03      DVILTENMFGDILSDEAAMLTGSLGMLPSASLGD-KFGLYEPVHGSAPDIAGQGIANPIA
Anc04      DVILTENMFGDILSDEAAMLTGSLGMLPSASLGD-GFGLYEPVHGSAPDIAGQGIANPIA
Anc05      DVILTENMFGDILSDEAAMLTGSLGMLPSASLGESGFGLYEPAGGSAPDIAGQGIANPIA
Anc06      DVILCENMFGDILSDEAAMLTGSLGMLPSASLAESGFGLYEPAGGSAPDIAGQGIANPIA
Anc07      DVILCENMFGDILSDEAAMLTGSLGMLPSASLAESGFGLYEPAGGSAPDIAGQGIANPIA
Anc08      DVMLCSNLFGDILSDECAMLTGSMGMLPSASLAESGFGLYEPAGGSAPDIAGKGIANPIA
Anc09      DVMLCSNLFGDILSDECAMLTGSMGMLPSASLNESGFGLYEPAGGSAPDIAGKGIANPIA
Anc10      DVMLCSNLFGDILSDECAMITGSMGMLPSASLNESGFGLYEPAGGSAPDIAGKNIANPIA
Anc11      DVMLCSNLFGDILSDECAMITGSMGMLPSASLNESGFGLYEPAGGSAPDIAGKNIANPIA
EcIPMDH    DVLLCSNLFGDILSDECAMITGSMGMLPSASLNEQGFGLYEPAGGSAPDIAGKNIANPIA
```

```
         298                                                        357
Anc01      AILSAAMMLRYSFGMEEAAEAIEQAVEKVLAEGYRTADIAQGGSKLVSTKEMGDAIVERL
Anc02      AILSAAMMLRYSFGMEEAAEAIERAVEKALAEGYRTADIAQGGSKLVSTKEMGDAIVERL
Anc03      AILSAAMMLRYSFGMEEAAEAIEKAVEKALAEGYRTADIAQGGSKLVSTKEMGDAIVERL
Anc04      AILSAAMMLRYSFGMEEAAEAIEKAVEKALAEGYRTADIAQGGSKKVSTKEMGDAIVERL
Anc05      QILSAAMMLRYSFGMEEAAQAIEKAVEKVLAEGYRTADIAQDGSKKVSTKEMGDAIVERL
Anc06      QILSAAMMLRYSFGMEEAAQAIEKAVEKVLAEGYRTADIAQEGSTKVSTKEMGDAIVEAL
Anc07      QILSAAMMLRYSFGMEEAAQAIERAVEKVLAQGYRTGDIAQEGSTKVSTKEMGDAIVEAL
Anc08      QILSAALMLRYSFKQEEAAQAIERAVEKVLAQGYRTGDIAQEGSTKVSTTEMGDAIVEAL
Anc09      QILSAALMLRYSLKQEEAAQAIERAVSKALAQGYLTGDLAQQGHTAVSTTEMGDAIAEAV
Anc10      QILSAALMLRYSLKQEEAAQAIERAVSKALAQGYLTGDLAQ-GHTAVSTDEMGDAIAEAV
Anc11      QILSAALMLRYSLKQEEAAQAIERAVSKALAEGYLTGDLAG-GHAAVSTSEMGDAIASYV
EcIPMDH    QILSLALLLRYSLDADDAACAIERAINRALEEGIRTGDLAR-GAAAVSTDEMGDIIARYV
```

```
         358
Anc01      ESRV
Anc02      ESSV
Anc03      ESSV
Anc04      EES-
Anc05      EES-
Anc06      EES-
Anc07      KES-
Anc08      KES-
Anc09      KEGV
Anc10      KEGV
Anc11      KEGV
EcIPMDH    AEGV
```

**Figure S2. A multiple amino acid sequence alignment of ancestral IPMDHs and EcIPMDH.** The positions targeted for site-directed mutagenesis are highlighted in color. The residue numbers above the sequences correspond to Anc05.

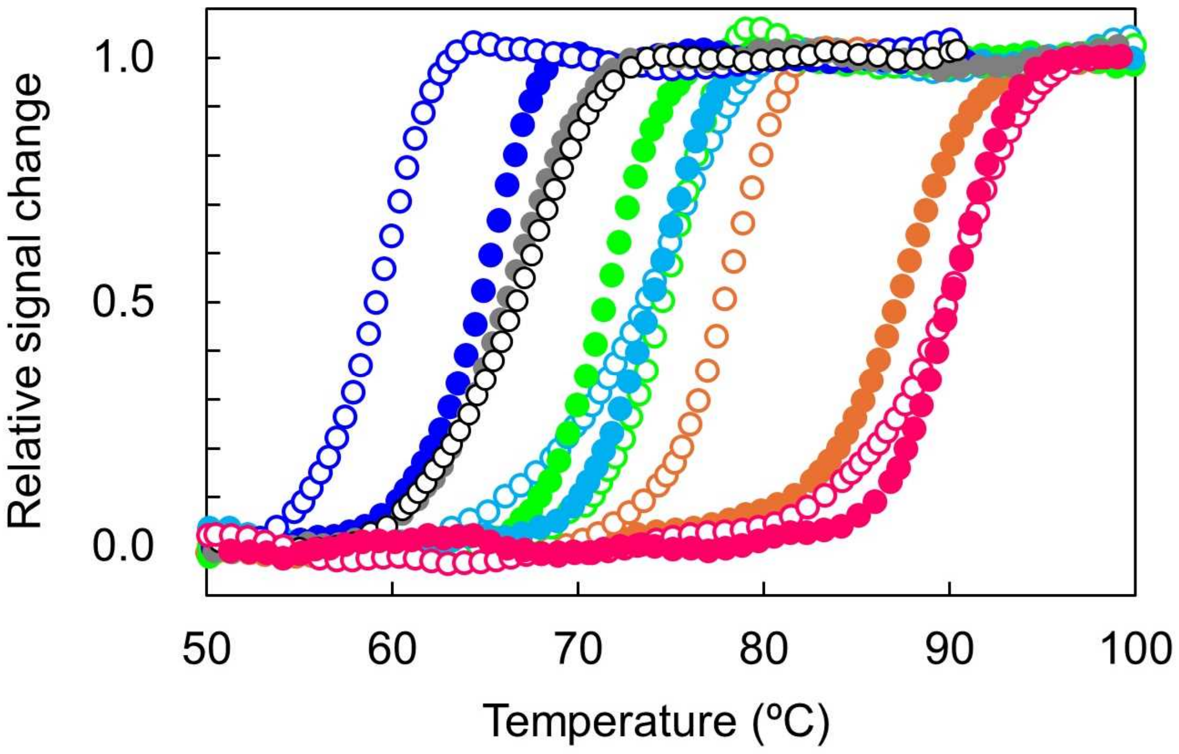


**Figure S3.** Temperature-induced unfolding curves of the ancestral and *E. coli* IPMDHs. The change in ellipticity at 222 nm was monitored as a function of temperature. The temperature was increased at a rate of 1.0°C/min. The samples comprised 5.0 μM protein in 20 mM potassium phosphate (pH 7.6), 0.5 mM EDTA. Each experiment was performed in duplicate with identical unfolding curves within experimental error. The plots were normalized with respect to the baseline of the native and denatured states. Magenta closed circles, Anc01; magenta open circles, Anc02; orange closed circles, Anc03; orange open circles, Anc04; green closed circles, Anc05; green open circles, Anc06; light blue closed circles, Anc07; light blue open circles, Anc08; dark blue closed circles, Anc09; dark blue open circles, Anc10; gray closed circles, Anc11; black open circles, EcIPMDH.

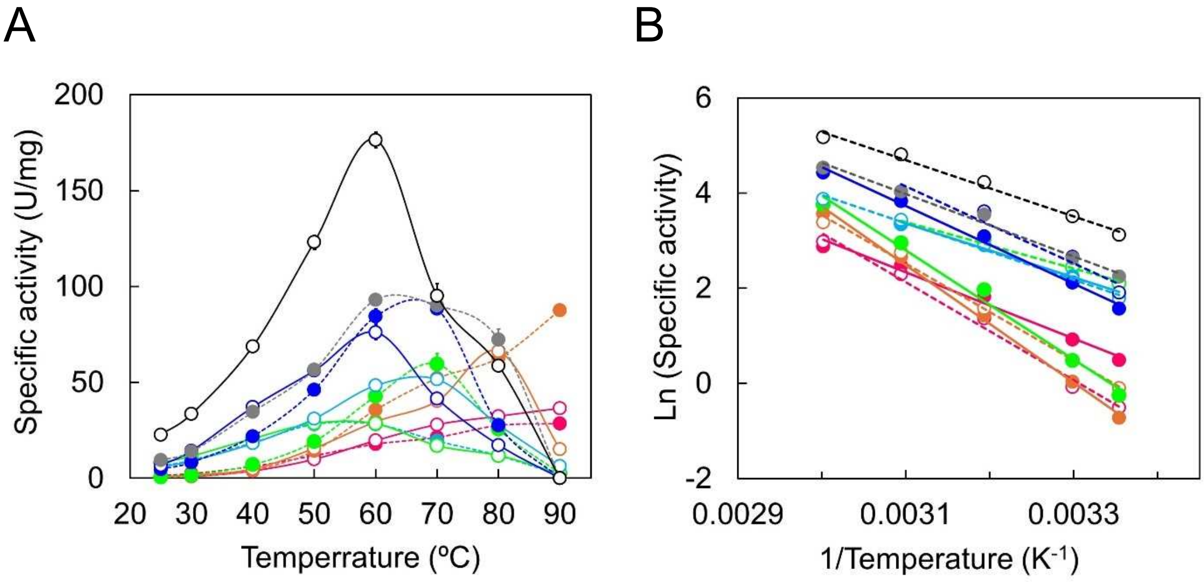


**Figure S4. Specific activities of the ancestral and *E. coli* IPMDHs.** (**A**) Plot of specific activity as a function of temperature. The assay solution was composed of 50 mM HEPES (pH 8.0), 100 mM KCl, 5 mM $MgCl_2$, 0.2 mM D-3-IPM, 5 mM $NAD^+$, and 0.3–3.0 μM protein. Each value is the average of three measurements. (**B**) Arrhenius plot of the specific activities of the ancestral and *E. coli* IPMDHs. Magenta closed circles and solid lines, Anc01; magenta open circles and dashed lines, Anc02; orange closed circles and solid lines, Anc03; orange open circles and dashed lines, Anc04; green closed circles and solid lines, Anc05; green open circles and dashed lines, Anc06; light blue closed circles and solid lines, Anc07; light blue open circles and dashed lines, Anc08; dark blue closed circles and solid lines, Anc09; dark blue open circles and dashed lines, Anc10; gray closed circles and solid lines, Anc11; black open circles and dashed lines, EcIPMDH.

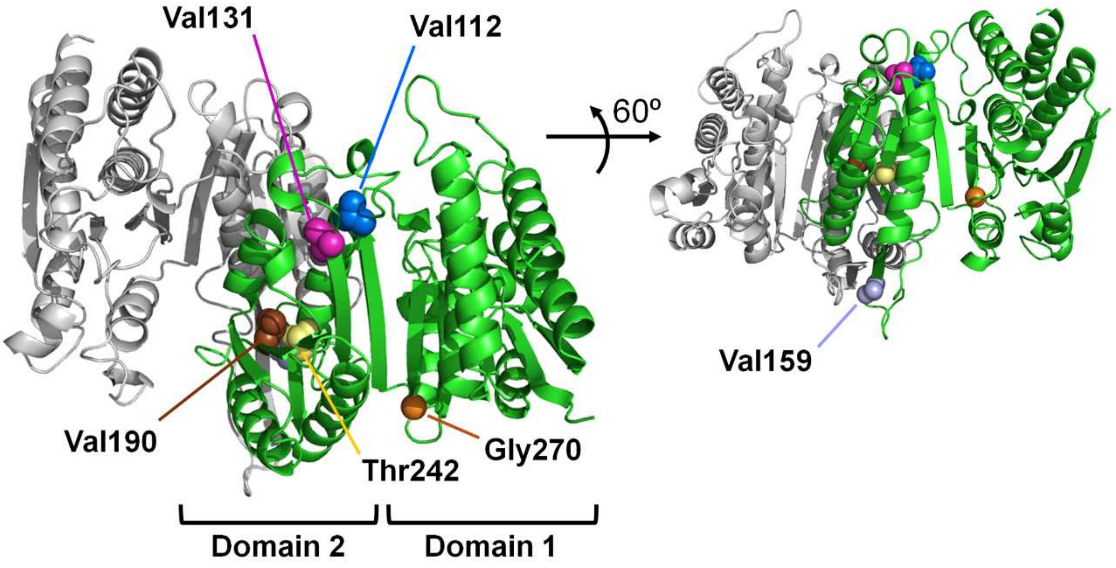


**Figure S5. A modelled dimeric structure of Anc05 predicted using Alphafold2 (37).** The subunits of the proteins are each colored differently. Each subunit is divided into two structural domains; domain 1 includes the N- and C-termini, and domain 2 includes the subunit interface. Residues at positions 112, 131, 15, 190, 242, and 270 in each subunit are highlighted. The side chains of residues 112 and 131 interact with each other, and the side chains of residues 190 and 242 also interact. Residues 112 and 270 are each located at a terminus of a β-strand connecting the two domains.

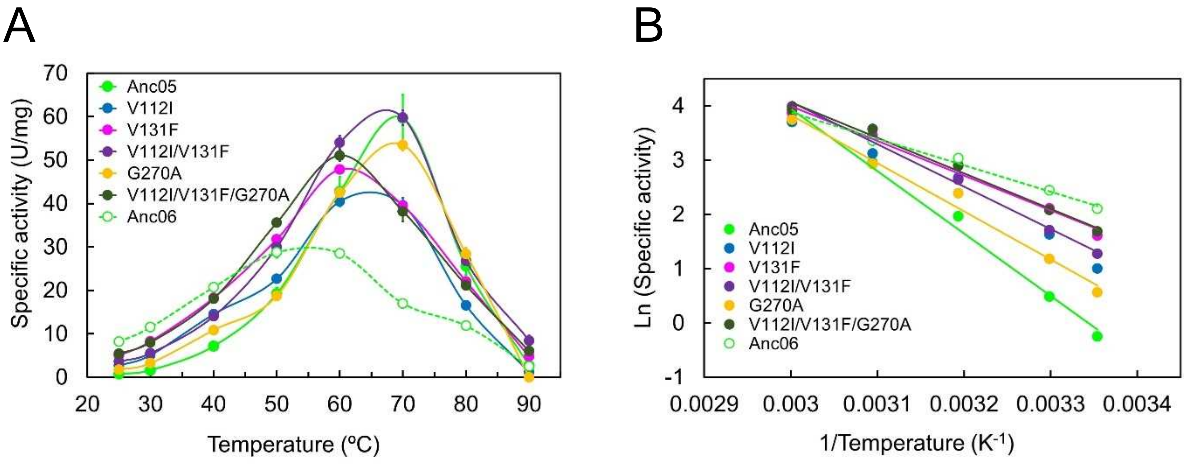


**Figure S6. Specific activities of Anc05, its mutants, and Anc06.** (**A**) Plot of specific activity as a function of temperature. The assay solution was composed of 50 mM HEPES (pH 8.0), 100 mM KCl, 5 mM $MgCl_2$, 0.2 mM D-3-IPM, 5 mM $NAD^+$, and 0.3–3.0 μM protein. Each value is the average of three measurements. (**B**) Arrhenius plot of the specific activities of ancestral IPMDHs and the mutants of Anc05.

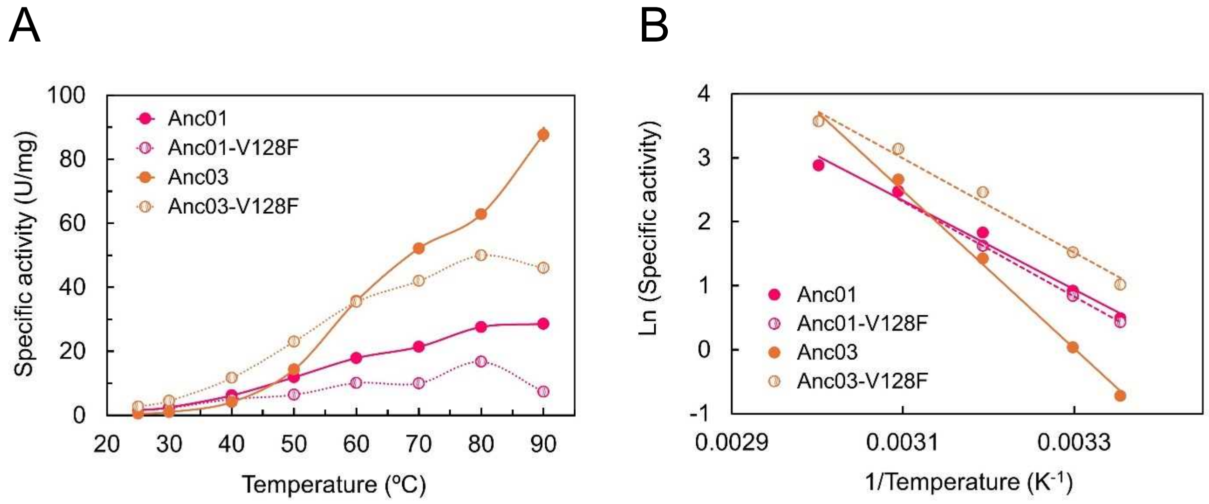


**Figure S7. Specific activities of Anc01, Anc03, and their V128F mutants.** (**A**) Plot of specific activity as a function of temperature. The assay solution was composed of 50 mM HEPES (pH 8.0), 100 mM KCl, 5 mM $MgCl_2$, 0.2 mM D-3-IPM, 5 mM $NAD^+$, and 0.3–3.0 μM protein. Each value is the average of three measurements. (**B**) Arrhenius plot of the specific activities of the ancestral IPMDHs and their mutants.

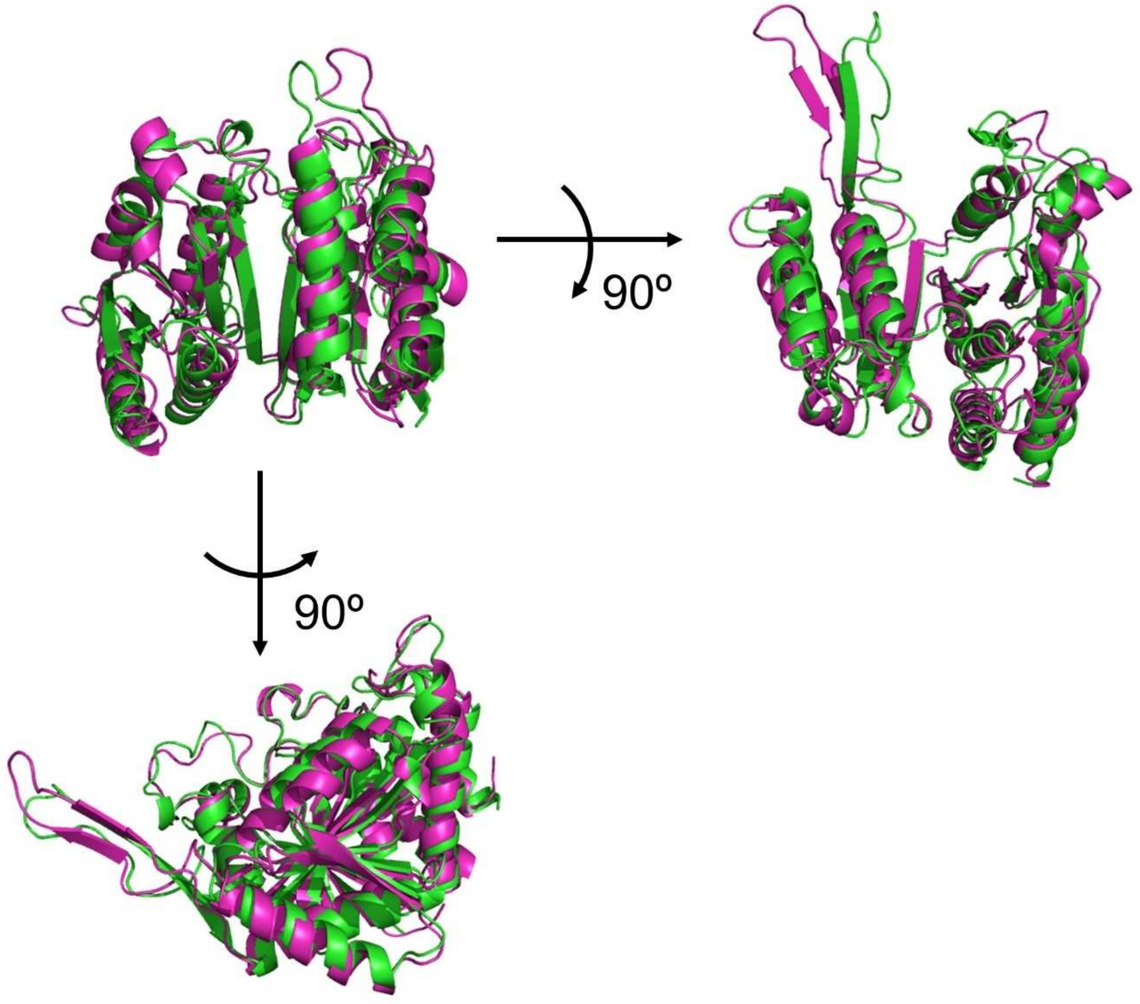


**Figure S8. Comparison of the modeled structure of Anc05 (colored green) predicted using Alphafold2 (37) and the crystal structure of EcIPMDH (colored magenta; PDB code 1cm7).** Superimposition was performed with PDBeFold ver. 2.59 (https://www.ebi.ac.uk/msd-srv/ssm/) and the superimposed structures were visualized with PyMOL (https://pymol.org). The two structures were reasonably similar with the average value of the $C_a$ root-mean-square deviations being 1.92 Å for 342 aligned residues.

**Table S1. The numerical data for $T_m$, $E_a$, $k_{cat}$ , $K_m^{D\text{-}3\text{-}IPM}$, and $K_m^{NAD}$ for ancestral IPMDHs, their mutants, and EcIPMDH.**

| | $T_m$ [a] (°C) | $E_a$ (kJ/mol) | $k_{cat}$ [b] ($s^{-1}$) | $K_m^{D\text{-}3\text{-}IPM}$ [b] (μM) | $K_m^{NAD}$ [b] (μM) |
|---|---|---|---|---|---|
| Anc01 | 90 | 58 | 1.0 ± 0.0 | 3.0 ± 0.2 | 56 ± 7 |
| Anc02 | 90 | 86 | 0.38 ± 0.01 | 1.7 ± 0.5 | 22 ± 2 |
| Anc03 | 87 | 103 | 0.33 ± 0.01 | 1.4 ± 0.3 | 12 ± 1 |
| V128F | 85 | 61 | 1.5 ± 0.1 | 3.0 ± 0.4 | 91 ± 23 |
| Anc04 | 78 | 85 | 0.70 ± 0.0 | 2.7 ± 0.8 | 27 ± 5 |
| Anc05 | 71 | 96 | 0.59 ± 0.03 | 1.7 ± 0.5 | 7.0 ± 1.6 |
| V112I | n.d. [c] | 62 | 1.8 ± 0.1 | 1.6 ± 0.4 | 25 ± 3 |
| V131F | n.d. [c] | 53 | 4.7 ± 0.1 | 3.0 ± 0.4 | 170 ± 10 |
| G270A | n.d. [c] | 74 | 0.90 ± 0.03 | 4.3 ± 1.1 | 10 ± 1 |
| V121I/V131F | n.d. [c] | 65 | 3.4 ± 0.1 | 1.8 ± 0.4 | 63 ± 5 |
| V121I/V131F/G270A | n.d. [c] | 55 | 3.8 ± 0.2 | 2.8 ± 0.4 | 280 ± 40 |
| Anc06 | 75 | 41 | 7.3 ± 0.5 | 2.5 ± 0.8 | 1700 ± 300 |
| Anc07 | 74 | 47 | 5.4 ± 0.3 | 2.5 ± 0.4 | 1800 ± 300 |
| Anc08 | 74 | 49 | 4.4 ± 0.1 | 2.2 ± 0.2 | 230 ± 30 |
| Anc09 | 64 | 68 | 4.0 ± 0.0 | 2.1 ± 0.4 | 11 ± 1 |
| Anc10 | 49 | 67 | 3.9 ± 0.1 | 1.9 ± 0.1 | 27 ± 3 |
| Anc11 | 66 | 54 | 8.5 ± 0.4 | 1.8 ± 0.2 | 33 ± 7 |
| EcIPMDH | 67 | 49 | 14 ± 0.5 | 3.7 ± 1.4 | 57 ± 9 |

[a] The $T_m$ values were estimated from the data shown in Fig. S3 and Fig. 5C.

[b] $K_m$s, $k_{cat}$, and standard errors were calculated by nonlinear least-square fitting of the steady-state kinetic data to the Michaelis-Menten equation using the Enzyme Kinetics module of SigmaPlot Ver. 14.5 (Systat Software).

[c] $T_m$ values are not determined for the Anc05 mutants.

**Table S2. PCR oligonucleotide primers used for mutagenesis.**

| Primer | Sequence (5'→3') [a] |
|---|---|
| T7_forward [b] | TAATACGACTCACTATAGG |
| T7_reverse [b] | GCTAGTTATTGCTCAGCGG |
| Anc05_V112I_forward | GCCCGGCCAAAATCTATCCGGCG |
| Anc05_V112I_reverse | CGCCGGATAGATTTTGGCCGGGC |
| Anc05_V131F_forward | GTTGGAGAAGGTTTCGACATTCTGGTAG |
| Anc05_V131F_ reverse | CTACCAGAATGTCGAAACCTTCTCCAAC |
| Anc05_V159F_forward | CGAAGAGAAAGCATTTGACACGATGG |
| Anc05_V159F_reverse | CCATCGTGTCAAATGCTTTCTCTTCG |
| Anc05_V190I_forward | GAAAGTAACCTCAATCGACAAAGCAAATG |
| Anc05_V190I_reverse | CATTTGCTTTGTCGATTGAGGTTACTTTC |
| Anc05_T242C_forward | GACGTTATCCTGTGCGAAAACATGTTC |
| Anc05_T242C_ reverse | GAACATGTTTTCGCACAGGATAACGTC |
| Anc05_G270A_forward | GTGCCTCACTGGCTGAATCTGGGTTTG |
| Anc05_G270A_reverse | CAAACCCAGATTCAGCCAGTGAGGCAC |
| Anc01_V128F_forward | GTCGTAGAAGGGTTCGATATCCTTGTTG |
| Anc01_V128F_reverse | CAACAAGGATATCGAACCCTTCTACGAC |
| Anc03_V128F_forward | GATCGTTGAAGGCTTCGATATCCTTGTTG |
| Anc03_V128F_reverse | CAACAAGGATATCGAAGCCTTCAACGATC |

[a] The mutated sequences are underlined.

[b] T7_forward and T7_reverse were used for all reactions.

**Data S1. Amino acid sequences of ancestral and *E. coli* IPMDHs in FASTA format.**

```
>Anc01
MTYKIAVLPGDGIGPEVVAEAVKVLEAVAEKFGLEFEFEEALIGGAAIDATGTPLPEETLEVCKESDA
VLLGAVGGPKWDNLPPDLRPERGLLKLRKALGLFANLRPAKVYPALVDASPLKPEVVEGVDILVVREL
TGGIYFGQPRGIDKGNERAVDTMVYTRSEIERIARVAFELARKRRKKVTSVDKANVLESSQLWREVVT
EVAKEYPDVELEHMYVDNCAMQLVRNPRQFDVILTENMFGDILSDEAAMLTGSLGMLPSASLGDKVGL
YEPVHGSAPDIAGQGIANPIAAILSAAMMLRYSFGMEEAAEAIEQAVEKVLAEGYRTADIAQGGSKLV
STKEMGDAIVERLESRV

>Anc02
MTYKIAVLPGDGIGPEVVAEAVKVLEAVAEKFGLEFEFEEALIGGAAIDAHGTPLPEETLEVCKESDA
VLLGAVGGPKWDNLPPDLRPERGLLKLRKALGLFANLRPAKVYPALVDASPLKPEVVEGVDILVVREL
TGGIYFGQPRGRDEGERAVDTMVYTRSEIERIARVAFEVARKRRKKVTSVDKANVLETSQLWREVVTE
VAKEYPDVELEHMYVDNCAMQLVRDPRQFDVILTENMFGDILSDEAAMLTGSLGMLPSASLGDKFGLY
EPVHGSAPDIAGQGIANPIAAILSAAMMLRYSFGMEEAAEAIERAVEKALAEGYRTADIAQGGSKLVS
TKEMGDAIVERLESSV

>Anc03
MTYKIAVLPGDGIGPEVVAEAVKVLEAVAEKFGLEFEFEEALIGGAAIDAHGHPLPEETLEVCKESDA
VLLGAVGGPKWDNLPPELRPERGALLPLRKALGLFANLRPAKVYPALVDASPLKPEIVEGVDILVVRE
LTGGIYFGQPRGRDEGERAVDTMVYTRSEIERIARVAFEVARKRRKKVTSVDKANVLETSQLWREVVT
EVAKEYPDVELEHMYVDNCAMQLVRDPRQFDVILTENMFGDILSDEAAMLTGSLGMLPSASLGDKFGL
YEPVHGSAPDIAGQGIANPIAAILSAAMMLRYSFGMEEAAEAIEKAVEKALAEGYRTADIAQGGSKLV
STKEMGDAIVERLESSV

>Anc04
MTYKIAVLPGDGIGPEVMAEALKVLEAVSKKFGLEFEFEEALVGGAAIDAHGHPLPEETLEVCEESDA
ILFGAVGGPKWENLPPEQQPERGALLPLRKAFGLFANLRPAKVFPALVDASPLKPEIVEGVDILVVRE
LTGGIYFGQPKGRDEGKEKAVDTMVYSRSEIERIARVAFEVARKRRKKVTSVDKANVLTTSVLWREVV
TEVAKDYPDVELEHMYVDNCAMQLVRDPRQFDVILTENMFGDILSDEAAMLTGSLGMLPSASLGDGFG
LYEPVHGSAPDIAGQGIANPIAAILSAAMMLRYSFGMEEAAEAIEKAVEKALAEGYRTADIAQGGSKK
VSTKEMGDAIVERLEES

>Anc05
MKTYKIAVLPGDGIGPEVMAEALKVLDAVSKKFGLEFEYEEAHVGGAAIDNHGTPLPEETLKICEESD
AILFGSVGGPKWENLPPEQQPERGALLPLRKHFGLFANLRPAKVYPALVDASPLKEEIVGEGVDILVV
RELTGGIYFGQPKGRDEGEEKAVDTMVYSRSEIERIARVAFEVARKRRKKVTSVDKANVLTTSVLWRE
VVTEVAKDYPDVELEHMYVDNAAMQLVRNPRQFDVILTENMFGDILSDEAAMLTGSLGMLPSASLGES
GFGLYEPAGGSAPDIAGQGIANPIAQILSAAMMLRYSFGMEEAAQAIEKAVEKVLAEGYRTADIAQDG
SKKVSTKEMGDAIVERLEES

>Anc06
MKTYKIAVLPGDGIGPEVMAEALKVLDAVSKKFGLTFEYEEAHVGGAAIDNHGSPLPEETLKICEESD
AILFGSVGGPKWENLPPEQQPERGALLPLRKHFGLFANLRPAKIFPALAHASPLKEEIVGEGFDILVV
RELTGGIYFGQPKGREGEGEEEKAFDTMVYSRSEIERIARVAFEVARKRRKKVTSIDKANVLTTSVLW
REVVTEVAKDYPDVELEHMYVDNAAMQLVRNPRQFDVILCENMFGDILSDEAAMLTGSLGMLPSASLA
ESGFGLYEPAGGSAPDIAGQGIANPIAQILSAAMMLRYSFGMEEAAQAIEKAVEKVLAEGYRTADIAQ
EGSTKVSTKEMGDAIVEALEES
```

>Anc07
MKTYKIAVLPGDGIGPEVMAEALKVLDAVSKKFGLTFEYEEAHVGGAAIDNHGSPLPEETLKICEESD
AILFGSVGGPKWENLPPEQQPERGALLPLRKHFGLFANLRPAKIFPALAHASPLKEEIVGEGFDILVV
RELTGGIYFGQPKGREGEGEEEKAFDTMVYSRSEIERIARVAFEAARKRRKKVTSIDKANVLTTSVLW
REVVTEVAKDYPDVELEHMYVDNAAMQLVRNPRQFDVILCENMFGDILSDEAAMLTGSLGMLPSASLA
ESGFGLYEPAGGSAPDIAGQGIANPIAQILSAAMMLRYSFGMEEAAQAIERAVEKVLAQGYRTGDIAQ
EGSTKVSTKEMGDAIVEALKES

>Anc08
MTKTYKIAVLPGDGIGPEVMAEALKVLDAVSKKFGLNFEYEEANVGGAAIDNHGSPLPESTLKLCEES
DAILFGSVGGPKWEHLPPNQQPERGALLPLRKHFKLFCNLRPAKIYQALAHASPLRADIVGEGFDILV
VRELTGGIYFGQPKGREGEGEEEKAFDTMVYSRSEIERIARIAFEAARLRRKKVTSIDKANVLTTSVL
WREVVTEVAKDYPDVELEHMYVDNAAMQLVKNPRQFDVMLCSNLFGDILSDECAMLTGSMGMLPSASL
AESGFGLYEPAGGSAPDIAGKGIANPIAQILSAALMLRYSFKQEEAAQAIERAVEKVLAQGYRTGDIA
QEGSTKVSTTEMGDAIVEALKES

>Anc09
MTKTYKIAVLPGDGIGPEVMAEALKVLDAVEQKFGLNFEYSEYDVGGAAIDNHGCPLPESTLKACEES
DAILFGSVGGPKWEHLPPNQQPERGALLPLRKHFKLFCNLRPAKIYQGLEHASPLRADISAKGFDILC
VRELTGGIYFGQPKGREGEGEHEKAFDTMVYSRREIERIARIAFEAARLRRKKVTSIDKANVLASSVL
WREVVTEVAKDYPDVELEHMYIDNATMQLVKNPSQFDVMLCSNLFGDILSDECAMLTGSMGMLPSASL
NESGFGLYEPAGGSAPDIAGKGIANPIAQILSAALMLRYSLKQEEAAQAIERAVSKALAQGYLTGDLA
QQGHTAVSTTEMGDAIAEAVKEGV

>Anc10
MTKTYKIAVLPGDGIGPEVMAEALKVLDAVEQKFGLNFEYSEYDVGGAAIDNHGCPLPESTLKACEES
DAILFGSVGGPKWEHLPPNQQPERGALLPLRKHFKLFCNLRPAKIYQGLEHASPLRADISAKGFDILC
VRELTGGIYFGQPKGREGEGEHEKAFDTEVYSRREIERIARIAFESARLRRKKVTSIDKANVLASSIL
WREVVTEVAKDYPDVELEHMYIDNATMQLVKNPSQFDVMLCSNLFGDILSDECAMITGSMGMLPSASL
NESGFGLYEPAGGSAPDIAGKNIANPIAQILSAALMLRYSLKQEEAAQAIERAVSKALAQGYLTGDLA
QGHTAVSTDEMGDAIAEAVKEGV

>Anc11
MTKTYKIAVLPGDGIGPEVMAEALKVLDAVEQKFGLRFTYSEYDVGGAAIDNHGCPLPESTLKACEEA
DAVLFGSVGGPKWEHLPPNQQPERGALLPLRKHFKLFCNLRPAKIYQGLEHFSPLRADISAKGFDILC
VRELTGGIYFGQPKGREGEGEHEKAFDTEVYHRFEIERIARIAFESARLRRKKVTSIDKANVLQSSIL
WREVVTEVAKDYPDVELSHMYIDNATMQLIKDPSQFDVMLCSNLFGDILSDECAMITGSMGMLPSASL
NESGFGLYEPAGGSAPDIAGKNIANPIAQILSAALMLRYSLKQEEAAQAIERAVSKALAEGYLTGDLA
GGHAAVSTSEMGDAIASYVKEGV

>EcIPMDH
MSKNYHIAVLPGDGIGPEVMTQALKVLDAVRNRFAMRITTSHYDVGGAAIDNHGQPLPPATVEGCEQA
DAVLFGSVGGPKWEHLPPDQQPERGALLPLRKHFKLFSNLRPAKLYQGLEAFCPLRADIAANGFDILC
VRELTGGIYFGQPKGREGSGQYEKAFDTEVYHRFEIERIARIAFESARKRRHKVTSIDKANVLQSSIL
WREIVNEIATEYPDVELAHMYIDNATMQLIKDPSQFDVLLCSNLFGDILSDECAMITGSMGMLPSASL
NEQGFGLYEPAGGSAPDIAGKNIANPIAQILSLALLLRYSLDADDAACAIERAINRALEEGIRTGDLA
RGAAAVSTDEMGDIIARYVAEGV

**Data S2 (separate file). Specific activities and standard errors of IPMDHs presented as numerical values in Excel format.** Each specific activity represents the mean of three independent measurements.
